# Curcumin-mediated NRF2 induction limits inflammatory damage in preclinical models of cystic fibrosis

**DOI:** 10.1101/2024.03.17.585384

**Authors:** Stephen A Leon-Icaza, Maxence Fretaud, Sarahdja Cornélie, Charlotte Bureau, Laure Yatime, R Andres Floto, Stephen A Renshaw, Jean-Louis Herrmann, Christelle Langevin, Céline Cougoule, Audrey Bernut

**Affiliations:** Institute of Pharmacology and Structural Biology, University of Toulouse, CNRS, Toulouse, France; Université Paris-Saclay, INRAE, Université de Versailles St Quentin, Virologie et Immunologie Moléculaires, Jouy-en-Josas, France; Laboratory of Pathogens and Host Immunity, University of Montpellier, CNRS, Inserm, Montpellier, France; Molecular Immunity Unit, University of Cambridge Department of Medicine, MRC-Laboratory of Molecular Biology, Cambridge, UK; Cambridge Centre for Lung Infection, Royal Papworth Hospital, Cambridge, UK.; The Bateson Centre, School of Medicine and Population Health, University of Sheffield, Sheffield, UK; Université Paris-Saclay, Université de Versailles St Quentin, Inserm, Infection et Inflammation, Montigny-le-Bretonneux, France; Hôpital Raymond Poincaré, AP-HP, Groupe Hospitalo-universitaire Paris-Saclay, Garches, France.; Université Paris-Saclay, INRAE, Infectiologie Expérimentale des Rongeurs et des Poissons, Jouy-en-Josas, France.

**Author notes:** Equal contribution (Stephen A Leon-Icaza, Maxence Fretaud). **Author Contributions**: AB conceived the project, designed experiments, analysed data and wrote the manuscript with input from SAL-I, MF, LY, RAF, SAR, J-LH, CL and CC. LY performed structural analyses. SAL-I and CC were responsible for the *ex vivo* testing in organoid models. MF and CL performed SHG imaging. SC, CB and AB performed zebrafish experiments. CL, CC and AB guided and supervised the work. All authors contributed to the article and approved the submitted version.

**Keywords:** Cystic fibrosis, Curcumin, NRF2, Oxidative response, Neutrophilic inflammation, Tissue repair

## Abstract

Overactive inflammation is directly correlated with airway damage and early death in individuals with cystic fibrosis (CF), a genetic disorder caused by mutation in the *CFTR* gene. Reducing the impact of inflammatory damage is therefore a major concern in CF. Several studies indicate that a decrease in the nuclear factor erythroid 2-related factor-2 (NRF2) signaling in people with CF may hamper their ability to alleviate oxidative stress and inflammation, although the role of NRF2 in CF inflammatory damage has not been determined. Therefore, we examined whether the phytochemical curcumin, an activator of NRF2, might provide a beneficial effect in the context of CF.

Herein, combining *Cftr*-depleted zebrafish larvae as innovative biomedical model with CF patient-derived airway organoids (AOs), we sought to understand how NRF2 dysfunction leads to abnormal inflammatory status and impaired tissue remodeling, and determine the effects of curcumin in reducing inflammation and tissue damage in CF.

We demonstrate that NFR2 is instrumental in efficiently regulating inflammatory and repair processes *in vivo*, thereby preventing acute neutrophilic inflammation and tissue damage. Importantly, curcumin treatment restores NRF2 activity in both CF zebrafish and AOs. Curcumin reduces neutrophilic inflammation in CF context, by rebalancing the production of epithelial ROS and pro-inflammatory cytokines. Furthermore, curcumin alleviates CF-associated tissue remodeling and allows tissue repair to occur. Our findings demonstrate that curcumin reduces inflammatory damage by restoring normal NRF2 activity, since disruption of Nrf2 pathway abrogated the effect of treatment in CF zebrafish.

This work highlights the protective role of NRF2 in limiting inflammation and injury, and show that therapeutic strategies to normalize NRF2 activity using curcumin might simultaneously reduce inflammation and enhance tissue repair, and thus prevent infectious and inflammatory lung damage in CF.

## Introduction

Cystic fibrosis (CF), resulting from mutations in the gene encoding the CF transmembrane conductance regulator (CFTR) channel, is one of the commonest fatal genetic disorders in the world (1, 2). Pulmonary pathology is the prime cause of mortality and morbidity among individuals with CF and is therefore the main focus of therapeutic interventions.

In lungs, absence or functional failure of CFTR results in an abnormal airway surface environment predisposing the patients to airway obstruction, infections, inflammation and tissue remodeling (3). Inflammation is a natural response of host immunity to infection or injury. However, in CF, airway inflammation is both excessive and ineffective at clearing infectious burdens, causing progressive and irreversible lung injury, ultimately leading to pulmonary function impairment and premature death (4).

The characteristic feature of airway inflammation in CF is an early, disproportionate and sustained neutrophil-dominated phenotype (5). Whether inherent to the CFTR defect or in response to infections, a plethora of cellular and molecular mechanisms have been proposed to explain the onset of neutrophilic inflammation in CF, including the pathogenic role of epithelial oxidative response (6–8) and excessive production of pro-inflammatory cytokines. Interestingly, critical regulators of redox balance, such as the nuclear factor erythroid 2-related factor-2 (NRF2)-KEAP1 pathway, appear to be disrupted in several *in vitro* and *ex vivo* models of CF (9–12). NRF2 is a transcription factor playing a pivotal role in the lungs to mitigate oxidative and inflammatory response through the regulation of genes involved in oxidative stress, antioxidant defense and pro-inflammatory processes *via* the antioxidant response elements (AREs) signaling pathway (13). Although CF epithelia exhibit reduced NRF2 activity and have been associated with increased production of reactive oxygen species (ROS) and pro-inflammatory cytokines (9), it remains unclear whether the excessive neutrophilic inflammation in CF is related to defective NRF2 function.

In CF care, the use of conventional anti-inflammatory therapies, such as corticosteroids and ibuprofen, can sometimes be effective in limiting inflammatory damage, but their adverse effects have discouraged their long-term use and new treatment options are desired (14). Considering the protective role of NRF2, therapeutic strategies designed to activate NRF2 may have clinical benefit in CF by reducing oxidative stress, inflammation and subsequent cell damage in the lungs. Curcumin, widely known for its anti-inflammatory and antioxidant properties, is a potent NRF2 activator (15). However, the biological significance of curcumin treatment in CF-related inflammatory damage is not known.

Using *cftr*-depleted zebrafish (7, 16) as an innovative and relevant vertebrate model of CF, combined with CF patient-derived airway organoids (AOs) (10), we demonstrated that curcumin, by normalizing NRF2 activity, can alleviate neutrophilic inflammation in CF context by reducing oxidative response and the levels of pro-inflammatory cytokines. In addition, curcumin directly mitigates CF-associated tissue damage and allows tissue repair to occur. These findings serve as supportive evidence for future evaluation of curcumin in people with CF.

## Results

### Zebrafish and human NRF2 proteins are structurally conserved

Two putative co-orthologs of human NRF2 (hNRF2) have been described in zebrafish, Nrf2a and Nrf2b (17, 18). Sequence comparison revealed that Nrf2a resembles hNRF2 more closely than Nrf2b as it shares 46% sequence identity with the human protein and displays a similar domain organization, with seven Neh domains and highly conserved consensus motifs, in particular in the Neh1 (ARE-binding) and Neh2 (KEAP1-binding) domains (**Figures 1A; S1**). In contrast, Nrf2b diverges a bit more from the human protein, with only 32% sequence identity, long deletion regions and only six conserved Neh domains, the Neh4 domain being absent (**Figures S1; S2A**). Despite some degree of conservation, sequence identity remains medium between hNRF2 and the zebrafish Nrf2a and Nrf2b, which questions the true functional orthology between these proteins.

**Figure 1.**
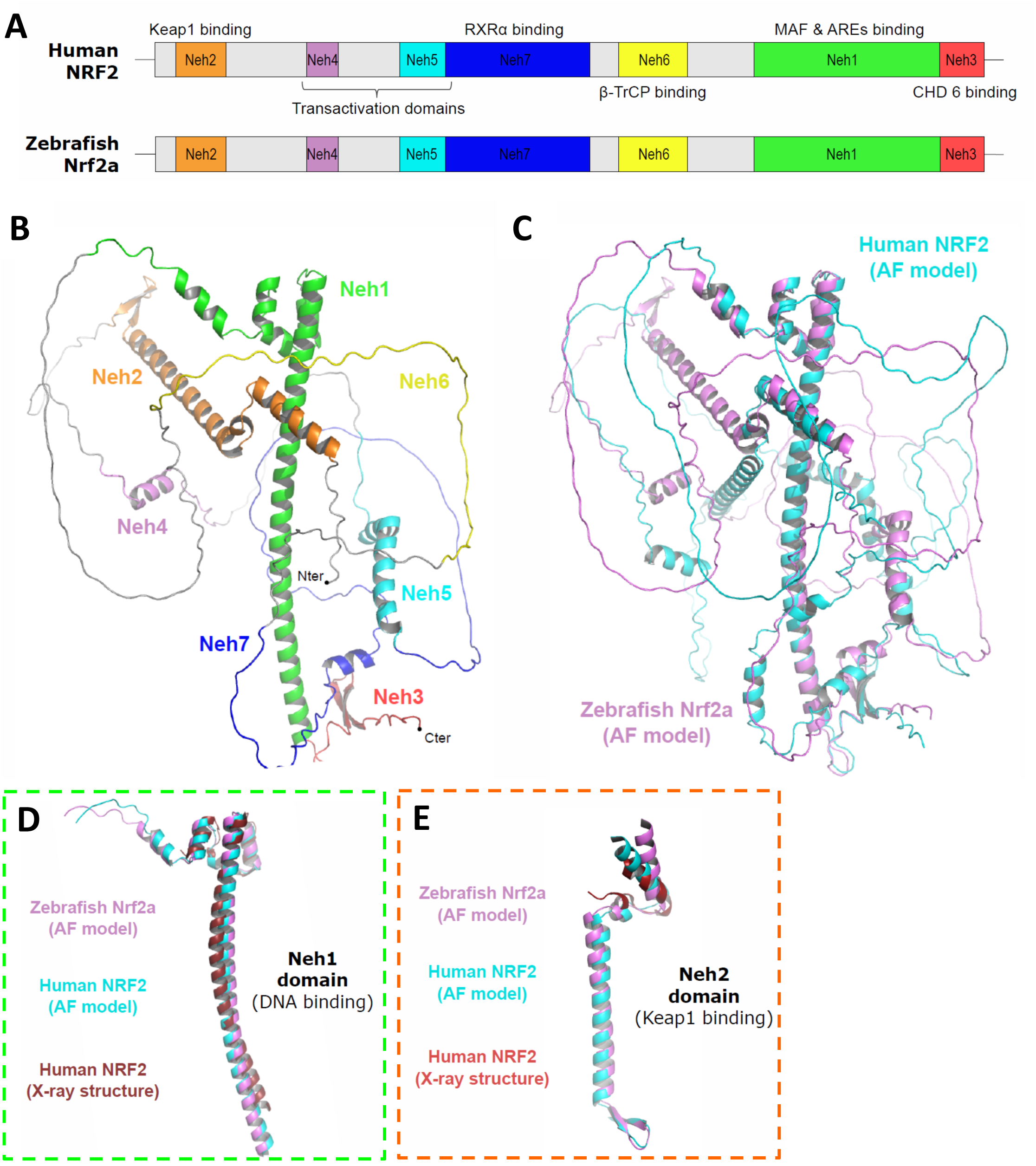
Structural analysis of zebrafish Nrf2a and comparison with human NRF2. (**A**) Domain organization of human NRF2 (hNRF2), with the target binding partners indicated for each domain, and predicted domain organization for zebrafish Nrf2a. All seven Neh domains seem to be present in Nrf2a. **(B)** 3D-structure of zebrafish Nrf2a predicted with AlphaFold. In this model, long unstructured regions alternate with shorter regions adopting defined secondary structures, mostly α-helices. The Neh domains are represented with the same colour code as in panel A. **(C)** Structural overlay of the AlphaFold-predicted model for zebrafish Nrf2a (light purple) with that of hNRF2 (cyan). The position of the secondary structure elements within the Nrf2a model matches well with those predicted for hNRF2. **(D)** Structural overlay of the Neh1 domains from AlphaFold-predicted Nrf2a structure, from AlphaFold-predicted hNRF2 structure, and from the X-ray structure of the human NRF2-MafG complex (red; PDB ID 7X5F; (20)). The Neh1 domain of Nrf2a align<u>s</u> well with the corresponding domains of hNRF2, with root mean square deviation (rmsd) values on Cα atoms ranging from 0.7 to 2.5 Å^2^. **(E)** Structural overlay of the Neh2 domains from AlphaFold-predicted Nrf2a structure, from AlphaFold-predicted hNRF2 structure, and from the X-ray structure of the hNRF2-Keap1 complex (red; PDB ID 3WN7; (21)). The Neh2 domain of Nrf2a aligns well with the corresponding domains of hNRF2, with rmsd values on Cα atoms ranging from 2.5 to 4.0 Å^2^.

To further address their resemblance, we performed structural analysis of the three proteins (hNFR2, Nrf2a and Nrf2b). As no experimental structure has been described for zebrafish Nrf2a yet, we used AlphaFold to predict its protein structure (19). As shown in **Figure 1B**, Nrf2a is proposed to adopt a largely unstructured 3D-architecture, alternating long coiled-coil regions with shorter, well-structured domains that correspond to the functional core of the Neh domains and are predicted with high confidence. In contrast, the low confidence of the prediction for coiled-coil regions suggests that they may fold differently when interacting with binding partners. Comparison of the AlphaFoldF model of Nrf2a with that of hNRF2 (**Figure 1C**) reveals a high structural similarity. In particular, the predicted secondary structure elements are placed at the same position in Nrf2a and hNRF2. Furthermore, the Neh1 and Neh2 domains of Nrf2a align well with the corresponding domains of hNRF2 from both the predicted AlphaFold model and available experimental structures of isolated domains (**Figures 1D-E**) (20, 21). A similar structural analysis was performed on Nrf2b and predicts a closely related structural organization although somewhat more divergent that hNRF2 (**Figures S2B-E**).

Taken together, these data suggest that zebrafish Nrf2 proteins, and in particular Nrf2a, are structural homologs of hNRF2, reinforcing the idea that they ensure similar biological functions, and thus validating the use of zebrafish model to study the function of NRF2 *in vivo*.

### CF-mediated NRF2 deficiency promotes overactive inflammation *in vivo*

To gain further insight into the impact of a dysfunctional CFTR on NRF2 activity *in vivo*, the expression of *nrf2a* and *nrf2b* in normal (control) and *cftr-*depleted (*cftr* morphant (MO)) zebrafish larvae was first evaluated following injury-induced inflammation (**Figure 2A**, (7)). While injury consistently triggered *nrf2a* response in control larvae, comparative RT-qPCR analyses revealed a significant downregulation of *nrf2a* level expression in CF animals (**Figure 2B**). Conversely, we found that *cftr* deficiency resulted in increased expression of *nrf2b* compared to the normal condition (**Figure 2B**).

**Figure 2.**
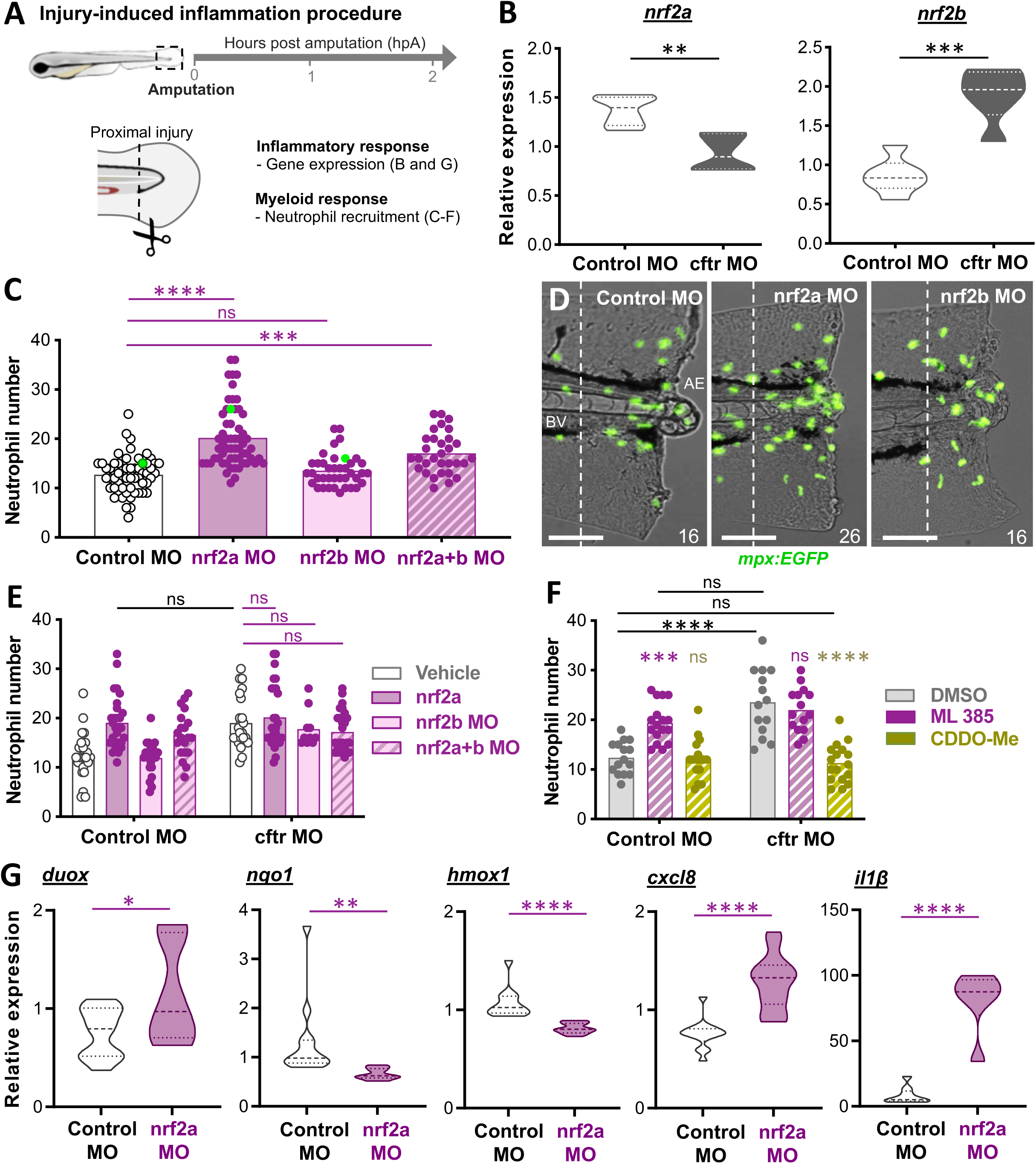
Injury-induced inflammatory responses are exacerbated in the absence of Nrf2 signaling in zebrafish larvae. (**A**) Schematic of aseptic injury-induced inflammation assay and analyses. Caudal fin amputations (dotted lined) were performed proximally through the notochord (N) at 3 days post-fertilization (dpf). **(B)** Control MO and *cftr* MO (generated using the validated *cftr* morpholino (16)) were caudal fin amputated then the level expression of mRNA for *nrf2a* (left) and *nrf2b* (right) genes are evaluated at 2 hours post-amputation (hpA). Graphs showing the fold change over total tissue from uninjured larvae (relative gene expression from at least 3 independent experiments performed in triplicates, mean and quartiles shown, Two-tailed Student *t*-test). **(C–F)** Neutrophil mobilization to tissue injury in the neutrophil-specific *TgBAC(mpx:EGFP)i114* line (22) is observed and enumerated at 2 hpA under a fluorescence microscope. The neutrophil count area is defined as the region between the blood vessels (BV) end and the amputation edge (AE). **(C)** Number of neutrophils mobilized to the wound in control MO, *nrf2a* MO, *nrf2b* MO or double *nrf2a* MO / *nrf2b* MO (*nrf2a+b* MO). Each dot represents the number of neutrophils at the wound in a single larva (3 independent experiments, One-way ANOVA with Dunnett’s comparisons test). **(D)** Representative images of injured tails (scale bars, 100 μm). The images are denoted as green data points in (C). **(E)** Inhibition of *nrf2a, nrf2b* or *nrf2a+b* was carried out in control and CF animals by injecting the *nrf2a* morpholino, the *nrf2b* morpholino or the *nrf2a* + *nrf2b* morpholinos reciprocally. Neutrophil recruitment assay (3 independent experiments, Two-way ANOVA with Tukey’s multiple comparisons test). **(F)** Control MO and *cftr* MO were treated with the NRF2 inhibitor ML 385, the NRF2 agonist CDDO-Me or DMSO (as mock control) prior to caudal fin amputation procedure, then injured and immediately put back in treatments. Neutrophil number at injured tails were enumerated at 2 hpA (3 independent experiments, Two-way ANOVA with Tukey’s multiple comparisons test). **(G)** Relative expression of mRNA for *duox, nqo1, hmox1, cxcl8 and il1β* at 2 hpA (*duox, cxcl8 and il1β*) or 8 hpA (*nqo1* and *hmox1*) from at least 3 independent experiments performed in triplicates (mean and quartiles shown, Two-tailed Student *t*-test).

Next, to determine how disrupted NRF2 activity might contribute to the hyper-inflammatory state in CF, expression of *nrf2a* and/or *nrf2b* were knocked-down in zebrafish using established morpholinos (18). As an exuberant influx of neutrophils is the hallmark of CF-related inflammation, we first investigated the consequence of Nrf2 inhibition on neutrophil response following injury by exploiting the *TgBAC(mpx:EGFP)i114* reporter line labeling neutrophils (22). Only zebrafish injected with the *nrf2a* morpholino were characterized by an increased recruitment of neutrophils to the wound compared to those injected with the control one, while no significant change was observed from animals injected with the *nrf2b* morpholino (**Figures 2C-D**). These results suggest that Nrf2a acts as a main regulator of neutrophilic inflammation in zebrafish, whereas Nrf2b shows no obvious function. To note, neither Nrf2a nor Nrf2b ablation had measurable impact on the total number of neutrophils (**Figures S3A-B**). Importantly, a similar increase in the number of wound-associated neutrophils was observed in *cftr*– and *nrf2a*-depleted larvae, and in the double *cftr/nrf2a*, *cftr/nrf2b* and *cftr/nrf2a+b*-depleted animals (**Figure 2E**). This is consistent with the hypothesis that NRF2 signaling is already disrupted in CF and might be involved in the excessive neutrophilic inflammation seen in this disease. We furthermore modulated NRF2 activity pharmacologically using the NRF2 blocker ML385 or the NRF2 activator bardoxolone methyl (CDDO-Me). Corroborating our finding using genetic approaches, we also observed that ML385 markedly exacerbated neutrophil response in wild-type animals (**Figure 2F**). In contrast, a protective effect of CDDO-Me exposure, acting to reduce neutrophilic inflammation, was demonstrated in CF larvae (**Figure 2F**). These findings suggest that NRF2 activation might alleviate excessive neutrophil responses in CF animals.

ROS generation by the dual oxidase DUOX is known to induce neutrophil chemoattraction (23). As shown in **Figures S4A-B**, pharmacological inhibition of ROS production reduced neutrophil response in *nrf2a*-depleted zebrafish, suggesting that exaggerated neutrophil mobilization associated with the loss of Nrf2a is due to a disrupted oxidative activity. To explore this hypothesis and to investigate the role of NRF2 in regulating redox balance, we next addressed whether Nrf2a ablation could affect oxidative responses in zebrafish. As expected, Nrf2a deficiency caused abnormal elevation of ROS production at the wound (**Figures S4C-D**), coinciding with increased expression of *duox* (**Figure 2G**). In contrast, *nqo1* and *hmox1*, two anti-oxidative NRF2 target genes, were downregulated in animals lacking Nrf2a (**Figure 2G**). Moreover, we found that loss of Nrf2a led to increased expression of pro-inflammatory genes *il8* and *il1β* (**Figures 2G; S4E-F**), two other important mediators of neutrophil chemotaxis (**Figure S4G**, (7)) known to be regulated by NRF2 (24, 25).

Collectively, these observations provide evidence that NRF2 activity is necessary to coordinate neutrophil trafficking. In particular, we show that *cftr*-depleted zebrafish recapitulate the abnormal NRF2 status associated with CF and support that dysfunctional NRF2 activity in CF airways contributes to neutrophilic inflammation, likely by causing deleterious changes in oxidative and pro-inflammatory responses.

### CF-mediated NRF2 deficiency alters tissue repair *in vivo*

Our previous work revealed that *cftr*-depleted zebrafish exhibit impaired tissue repair after injury (7). Considering the protective role of the NRF2 signaling pathway in repair processes (26), we next investigated whether dysfunctional NRF2 activity might impede tissue repair in CF. Since zebrafish caudal fins are known to undergo complete regeneration after amputation, we assessed host regenerative performance in the absence of Nrf2 signaling by quantifying fin regrowth post-amputation (**Figure 3A**, (7)). Fin regrowth was impaired in *nrf2a*-depleted zebrafish compared to controls, in association with a reduction in the area, length and width of regenerated fins (**Figures 3B-C**), whereas the loss of Nrf2b did not affect regrowth (**Figure S5**). These findings indicate that Nrf2a is the likely NRF2 paralogue that mediates tissue repair mechanisms in zebrafish. Importantly, similar regenerative defects were observed in both *cftr* and *nrf2a* MO, and in the double *cftr/nrf2a* MO (**Figure 3D**), suggesting that impaired tissue repair in CF zebrafish could be linked to defective Nrf2a. We next reasoned that excessive neutrophilic inflammation at wound (**Figure 2**) might contribute to the impaired tissue repair in *nrf2a*-depleted zebrafish. To address this question, we ablated neutrophils in zebrafish using the *csf3r* morpholino (27). As shown in **Figure S6**, removal of neutrophils significantly improved fin regrowth in *nrf2a* morphant animals.

**Figure 3.**
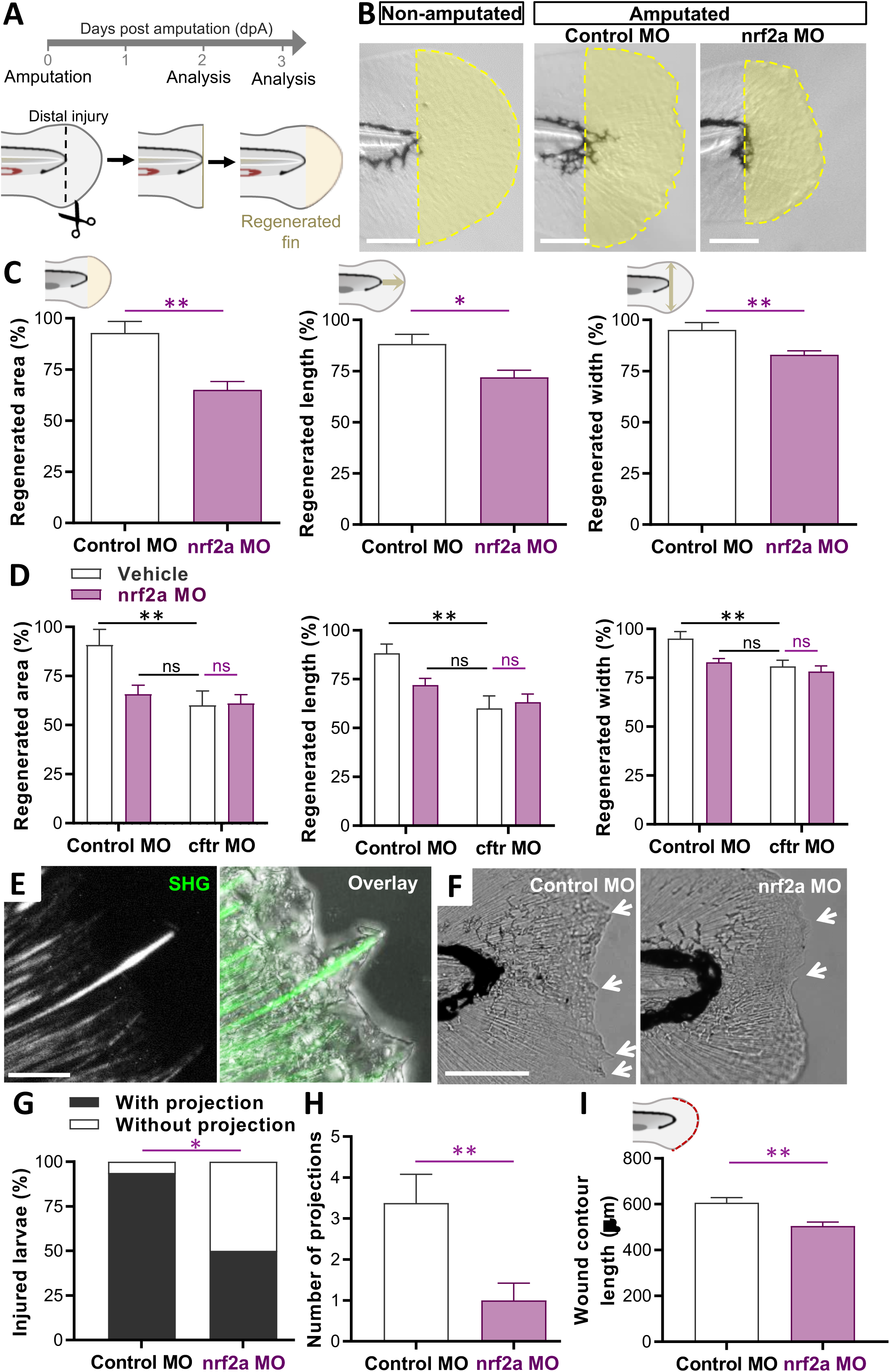
Nrf2a-depleted zebrafish exhibit reduced tissue repair responses after injury. (**A-I**) Tissue repair performance assessment. Caudal fin amputations were performed without injury to the notochord (dotted line) at 2 dpf, then the potential of tissue repair was evaluated at 2 **(E-I)** or 3 **(B-D)** days post-amputation (dpA). **(B-C)** Control MO and *nrf2a* MO were caudal fin amputated. Representative imaging of injured tail fin (scale bars, 100 μm) **(B)** and measurement of regenerated fin areas, lengths and widths **(C)** (n=14-18 from 3 independent experiments, Two-tailed Student *t*-test). Error bars represent standard error of the mean (SEM). The regenerated fin areas were defined as the region between the amputation edge and the end of the extended fin. The regenerated fin lengths were measured from the notochord tip to the end of the extended fin. The regenerated fin widths were defined as the max width measured from the amputation edge to the end of the extended fin. **(D)** Inhibition of *nrf2a* was carried out in WT and CF animals by injecting the *nrf2a* morpholino. Measurement of regenerated tail fin areas, lengths and widths in the presence or absence of *nrf2a* (n=16 from 3 independent experiments, Two-way ANOVA with Tukey’s multiple comparisons test). (**E**) Second harmonic generation (SHG) and high magnification bright field imaging showing epithelial projection associated with collagen fibers (scale bar, 10 μm). **(F)** Representative bright field microscopy of injured caudal fin showing extended collagen fiber-containing epithelial projections (arrows) pushing the healing plane forward (scale bar, 100 μm). Morpholino knockdown of *nrf2* expression reduced the appearance of projections, suggesting that Nrf2a modulates the formation of collagen fiber in zebrafish. **(G-H)** Proportion of larvae forming visible projections (**G**; n=14-16, Fisher test) and the mean ± SEM number of projections per larvae (**H**; n=14-16, Two-way ANOVA, Tukey’s multiple comparisons test) **(I)** Measurement of wound contour length in the presence or absence of *nrf2a* (n=14-18 from 3 independent experiments, Two-way ANOVA with Tukey’s multiple comparisons test).

It is known that NRF2 influences extracellular matrix (ECM) deposition and remodelling, including collagen formation (28), a critical step during tissue repair. In zebrafish, collagen projections at the wound edge guide epithelial growth during caudal fin regeneration and initiate tissue repair ((29, 30); **Figure 3E**). Since optimal tissue repair requires a functional NRF2 activity, we sought to characterize the effect of *nrf2* ablation on the formation of collagen projections during the process of regeneration. Using second harmonic generation (SHG) imaging to visualize collagen fibers, we found that the formation of collagen projections was disrupted in *nrf2a*-deficient larvae (**Figures 3F-I**). While the wound edge in control animals formed an uneven border containing collagen associated-epithelial projections, *nrf2a-*depleted larvae displayed a smooth wound edge and relative absence of such projections (**Figure 3F**). Following amputation, ≈ 90% of control larvae had visible projections while only ≈ 50% of *nrf2a-*depleted larvae displayed projections (**Figure 3G**). In addition, analyses revealed a reduced number of collagen projections in *nrf2a*-depleted animals compared to controls (**Figure 3H**). Finally, the contour of the wound edge was measured in control and *nrf2a* MO, as a marker of collagen fiber formation as well as a state of active tissue repair (29). The *nrf2a*-deficient larvae had a shorter contour length at the wound edge as compared to control larvae (**Figure 3I**). These findings indicate that Nrf2a deficiency impairs wound repair in zebrafish, at least in part through affecting the formation of collagen and epithelial projections.

Altogether, these findings provide further evidence that functional NRF2, by regulating collagen formation, is required for efficient tissue repair, and support the proposal that impaired NRF2 activity in CF airways could be involved in the abnormal tissue remodeling seen in this disease (31, 32).

### Curcumin-induced Nrf2 activation reduces CF-related inflammation in zebrafish

Current therapies targeting inflammation in CF have limited efficacy. Since NRF2 signaling represents an important mechanism in the regulation of inflammatory and tissue repair processes, it could be considered as a promising therapeutic target to prevent excessive inflammation and tissue damage in CF. We therefore examined whether turmeric-derived curcumin, a NRF2 activator, might provide a beneficial effect in the context of CF (**Figure 4A**).

**Figure 4.**
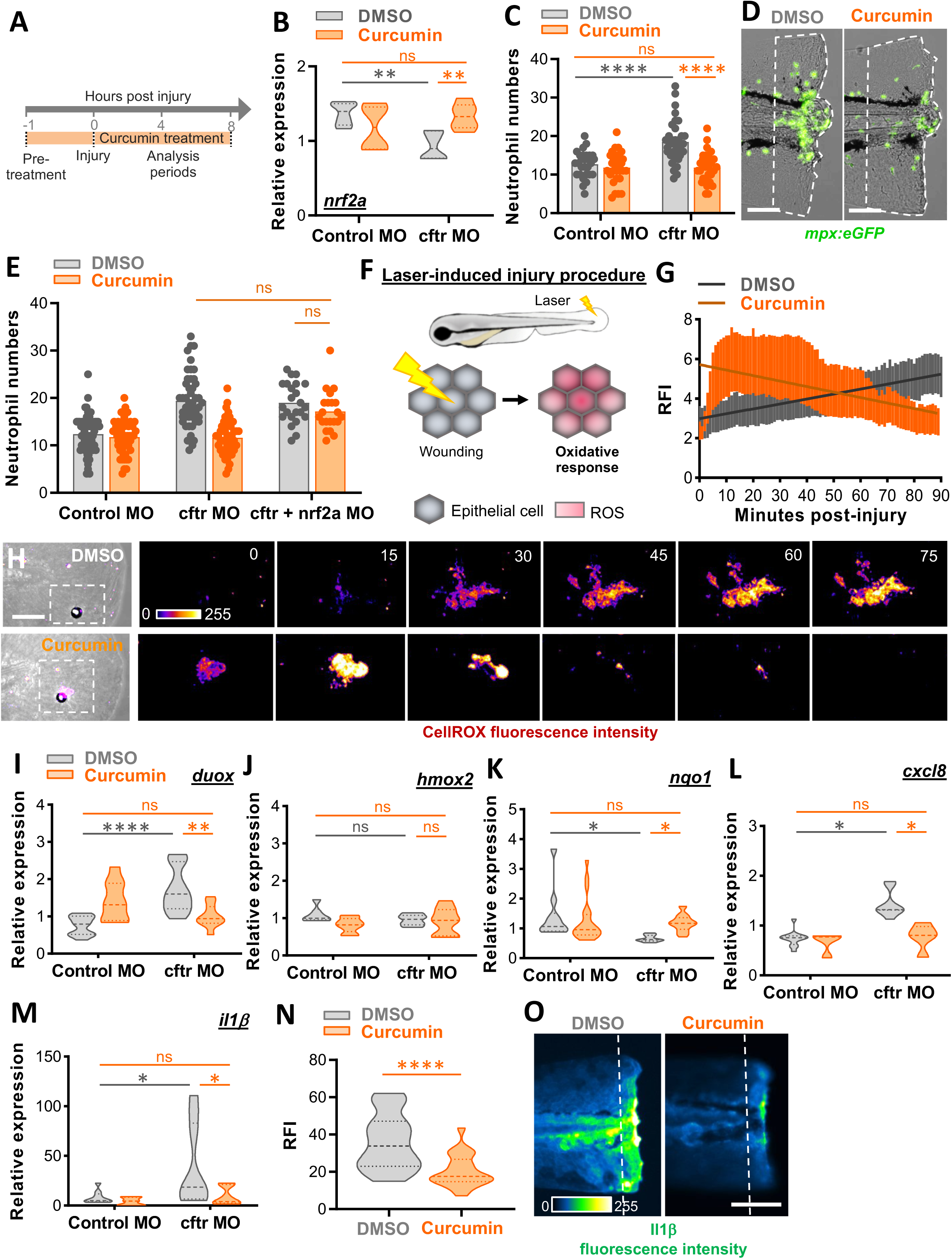
NRF2 activation by curcumin normalizes inflammation in *cftr*-depleted zebrafish. (**A**) Control MO and *cftr* MO were pre-treated with DMSO (as control) or curcumin before injury, then injured and immediately put back in treatments until analysis. **(B)** mRNA levels of *nrf2a* at 2 hpA (relative gene expression from at least 3 independent experiments performed in triplicates, Two-way ANOVA with Tukey’s multiple comparisons test). **(C)** Neutrophil number at the wound at 2 hpA (3 independent experiments, Two-way ANOVA with Tukey’s multiple comparisons test). **(D)** Representative images of amputation-induced neutrophil mobilization in *cftr*-MO treated or not with curcumin (scale bars, 100 μm). **(E)** Inhibition of *nrf2a* was carried out in CF animals by injecting the *nrf2a* morpholino. Neutrophil number at the wound at 2 hpA (3 independent experiments, Two-way ANOVA with Tukey’s multiple comparisons test). **(F)** Schematic illustration of laser-mediated injury assay triggering ROS production by epithelial cells. **(G-H)** *cftr* MO are stained with CellROX^®^, treated, laser-injured, then ROS production was time-lapse imaged and quantified over 90 minutes by confocal microscopy. **(G)** ROS intensity at the wound site over the time course of inflammation. Line of best fit shown is calculated by linear regression. *P-*value shown is for the difference between the 2 slopes (n=12, performed as 3 independent experiments). **(H)** Pseudocolored imaging of laser-injured caudal fin showing representative ROS production throughout inflammation. **(I-M)** mRNA levels of *duox, nqo1, hmox1, cxcl8 and il1β* at 2 hpA (*duox, cxcl8 and il1β*) or 8 hpA (*nqo1* and *hmox1*) (relative gene expression from at least 3 independent experiments performed in triplicates, Two-way ANOVA with Tukey’s multiple comparisons test). **(N-O)** *cftr* MO *Tg(il1b:eGFP-F)ump3* (72) were caudal fin amputated then the expression of *Il1β* was observed and analyzed. Relative *Il1β* intensity at 2hpA in *cftr* MO treated or not with curcumin (n=18 from 3 independent experiments; Mann-Whitney U test) **(N)** and associated pseudocolored photomicrographs of injured tails **(O)** revealing *il1β* expression at the wound-edge (scale bar, 100 μm).

Firstly, we confirmed that curcumin treatment was able to rescue disrupted *nrf2* expressions in Cftr-depleted zebrafish (**Figures 4B; S7**), and could promote Nrf2a activation. Curcumin efficiently reduced early neutrophil response to injury in CF larvae compared to DMSO-treated animals (**Figures 4C-D**; **S8**). Interestingly, despite the anti-inflammatory properties of curcumin, neutrophilic inflammation was similar in DMSO– and curcumin-treated larvae in the presence of normal Cftr activity (**Figures 4C; S8**). Additionally, since pulmonary infections are major features in CF (33), we were also interested in determining the effect of curcumin on neutrophilic inflammation in a context of infection. To do so, we used a range of CF-relevant bacteria, including *Staphylococcus aureus, Pseudomonas aeruginosa* and *Mycobacterium abscessus*, added into the media of injured larvae (**Figure S9A**). Microscopy observations revealed that neutrophil recruitment was significantly increased in response to the bacteria-infected injuries (**Figure S9B**). Having proven that curcumin does not alter bacterial growth *in vitro* (**Table 1**), we treated infected-CF zebrafish with curcumin and showed that the compound also alleviated hyper-neutrophilia in a context of infection (**Figure S9C**).

To validate the NRF2-targeted action of curcumin in our system, we knocked-down *nrf2a, nrf2b* or *nrf2a+b* expression in zebrafish and examined whether curcumin was still able to restore normal neutrophil responses in the absence of Nrf2 signaling. As shown in **Figure S10,** although wound-associated neutrophil number was significantly higher in *nrf2a*-depleted larvae than control animals when treated with curcumin, the treatment partially reduced neutrophilic inflammation in the absence of Nrf2a. However, when both *nrf2a* and *nrf2b* were depleted, curcumin failed to reduce neutrophil inflammation, supporting the notion that curcumin regulates neutrophil response through Nrf2-dependent mechanisms. Importantly, when *nrf2a* was depleted in CF animals, curcumin failed to reduce inflammation (**Figure 4E**), confirming that the presence of functional Nrf2 proteins, at least Nrf2a, is required for the anti-inflammatory action of curcumin to occur.

Excessive nuclear factor-κB (NF-κB) pathway activation has been described in the pathology of lung inflammation in CF (34), and was confirmed in zebrafish, with higher NFκB signal at wound in Cftr-depleted zebrafish compared to control animals (**Figures S11A-C**). In parallel of NRF2 activation, curcumin can directly inhibit NFκB signaling pathway (35). Therefore, the NFκB inhibitory action assigned to curcumin could also be considered. However, we found that curcumin exposure was not sufficient to reduced Nfκb signal in *cftr* MO (**Figure S11D**), suggesting that curcumin doesn’t reduce inflammation in CF through NFκB inhibition.

We next sought to understand the mechanisms by which curcumin-induced NRF2 activation reduces neutrophilic inflammation in CF animals. Since exaggerated production of ROS by epithelia contributes to the inflammatory pathogenesis in CF (36), we first proceeded to examine the potential benefits of curcumin in reducing oxidative stress in CF larvae. Unexpectedly, rapidly after laser-induced epithelial tissue injury, dynamic imaging and quantitative analyses revealed that curcumin-treated larvae exhibited a strong oxidative burst at the injury site (**Figures 4F-H**). However, while oxidative responses increased over time in DMSO-treated larvae, our observations indicated that curcumin gradually reduced ROS signal, suggesting that treatment might reduce ROS generation through the modulation of NRF2-induced genes. As expected, the anti-oxidative action of curcumin was confirmed by the downregulation of *duox* and the upregulation of *nqo1* gene in CF larvae treated with curcumin (**Figures 4I-K**), consistent with Nrf2 activation. Moreover, curcumin exposure ameliorated the increased Il8 and Il1β activity caused by CF, as demonstrated by qRT-PCR and/or microscopy analyses (**Figures 4L-O; S11E-G**).

Together these data indicate that curcumin efficiently alleviates CF-related neutrophilic inflammation, by reducing oxidative and pro-inflammatory responses, *via* NRF2 activation.

### Curcumin prevents tissue damage by improving tissue remodeling and repair in CF zebrafish

Pulmonary fibrosis and abnormal tissue remodeling disrupt lung function and cause death in CF. Considering the role of NRF2 pathway in repair processes (**Figure 3**), we next investigated whether curcumin-induced Nrf2a activation could rescue abnormal tissue remodeling and repair in CF zebrafish (7). As shown in **Figures 5A-C**, curcumin exposure significantly improved tissue repair in CF injured zebrafish. Furthermore, while the presence of bacteria worsened tissue damage and defective tissue repair phenotype (**Figures S12A-B**), curcumin was also able to enhance tissue repair in CF fish in the context of infection (**Figure S12C**).

**Figure 5.**
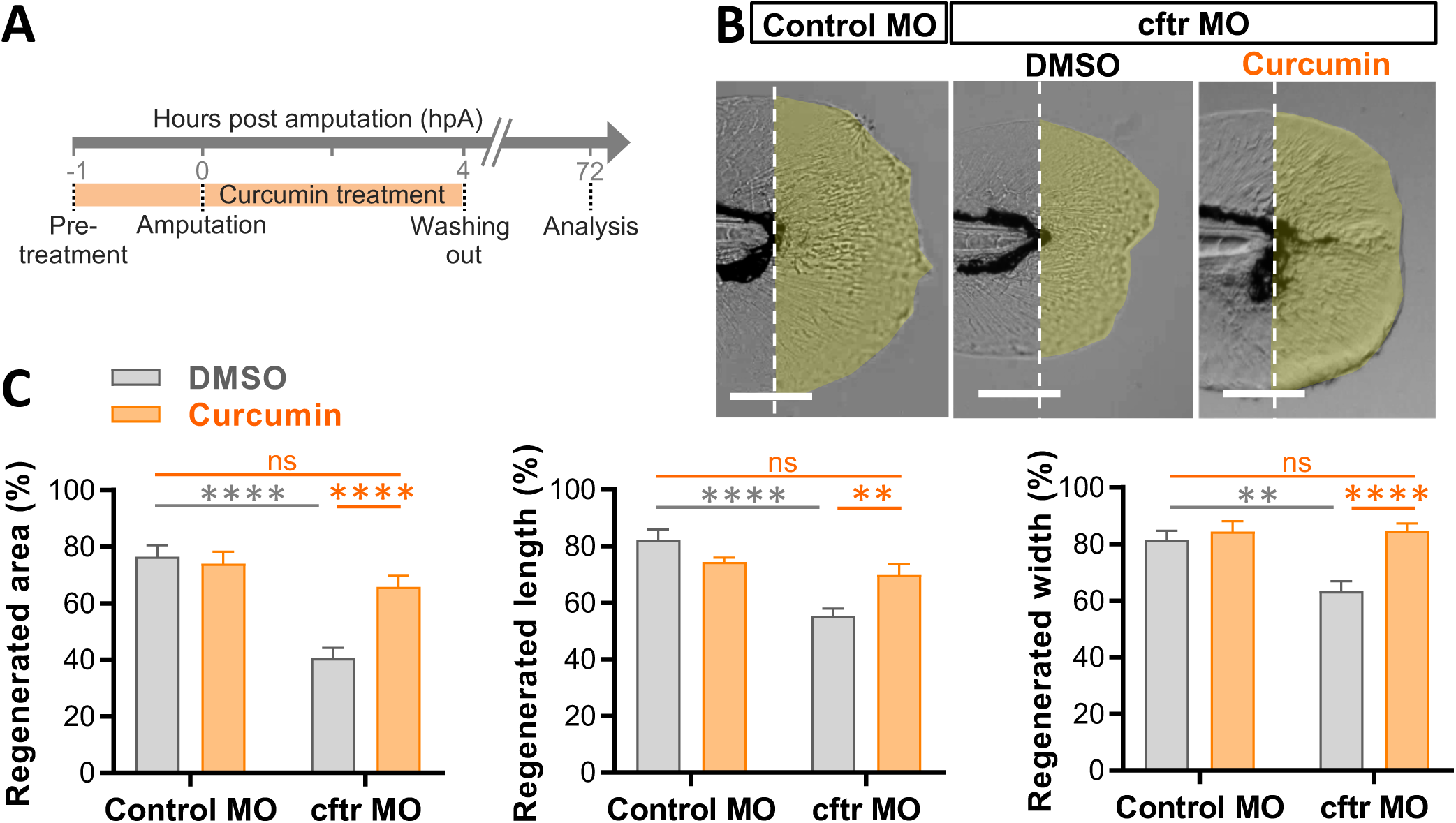
Curcumin improves defective tissue repair capacity in Cftr-depleted animals. (**A-C**) Control MO and *cftr* MO were treated with DMSO or curcumin prior to caudal fin amputation procedure, then injured and immediately put back in treatments for 4 h. The potential of tissue repair was evaluated at 72 hpA. **(B)** Brightfield microscopy of injured caudal fin showing tissue regeneration in *cftr* MO treated with curcumin *versus* DMSO (scale bars, 100 μm). **(C)** Measurement of regenerated caudal fin areas, lengths and widths following compounds exposure (n=20, Two-way ANOVA, Tukey’s multiple comparisons test).

Because proper collagen fiber production and reorganization are essential for functional tissue repair in zebrafish (30), we sought to further determine whether the deficit in caudal fin regeneration in CF zebrafish was associated with a defect in collagen remodeling. Firstly, comparative analysis of the control and Cftr-depleted zebrafish regenerated tissue revealed that collagen fibers did not project well in CF animals (**Figures 6A-D; S13A**). Furthermore, SHG imaging showed that collagen fibers were completely disrupted upon injury in CF animals, as measured by the area devoid of fibers (**Figures 6E-G; S13B**). Fiber alignment analysis (**Figures 6H-L**) indicated that this change correlates with a diminished number of fibers (**Figure 6I**) and a reduction in their lengths (**Figure 6J**). Importantly, a significant change in collagen fiber width was seen in cftr MO, with thicker fibers compared to control MO (**Figure 6K**), suggesting that Cftr influences injury-induced fiber thickening. Interestingly, this change was associated with an overexpression of the transforming growth factor-β (Tgf-β) (**Figure 6L**), a key pro-fibrotic cytokine / primary driver of fibrosis in CF (37). The population of fibers in CF fins also tended to be less perpendicular overall to the wound edge than their control counterparts (**Figure 6M**). Overall, regenerated tissues in CF animals consistently showed unaligned fibers compared to control, and were more prone to develop aberrant fiber extrusions into the intra-fiber space (**Figure 6N**). Interestingly, while mesenchymal cells are essential to guide the collagen fibers organization in zebrafish fins (38), we observed that disrupted fiber content and orientation coincided with an abnormal mesenchymal cell distribution and shape (**Figure S13C**). Remarkably, all these aberrant phenotypes are rescued by curcumin treatment (**Figures 6; S13**).

**Figure 6.**
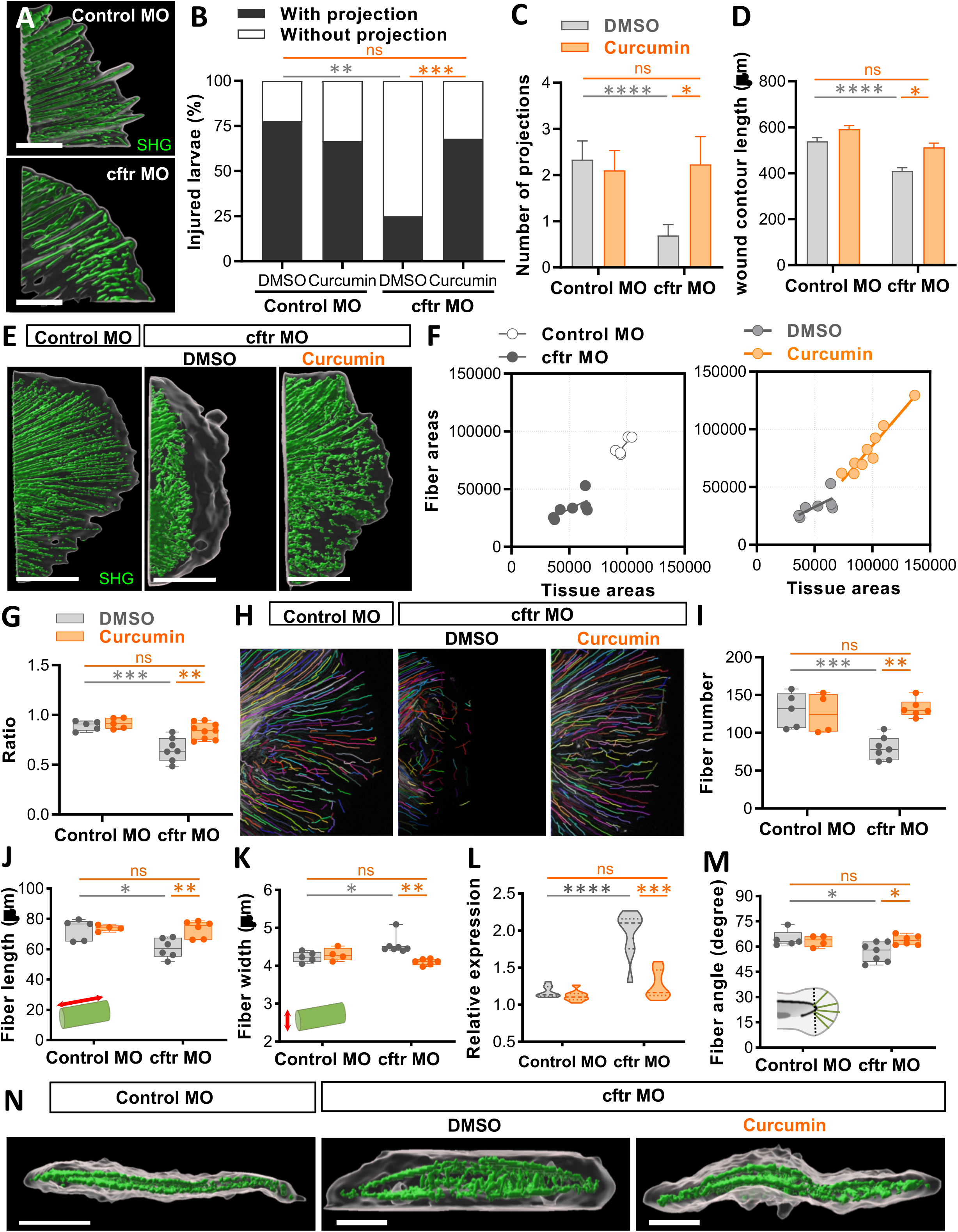
Curcumin reduces tissue damage by promoting remodeling of collagen fibers in CF zebrafish. Control MO and *cftr* MO were pre-treated with DMSO or curcumin prior to tail fin amputation procedure, then injured and immediately put back in treatments for 4 h. SHG microscopy of the regenerated tail fin was performed at 2 (**A-D**) or 3 dpA (**E-N**). **(A)** surface rendering of a 3D reconstruction showing the tissue wound edge in the bright field and corresponding SHG imaging Z projections to reveal collagen projections in injured tails in control *versus* CF animals (scale bars, 25 μm). **(B-C)** Proportion of larvae forming projections (**B**; n=20, Fisher test) and the number of projections per larvae (**C**; n=20, Two-way ANOVA, Tukey’s multiple comparisons test), in fishes treated or not with curcumin. **(D)** Measurement of wound contour length in fishes treated or not with curcumin (n=20 from 3 independent experiments, Two-way ANOVA with Tukey’s multiple comparisons test). **(E)** 3D surface-rendered reconstruction of collagen fibers in regenerated tails to illustrate the spatial organization of fiber relative to regenerated tissue with tissue wound edge in the bright field and corresponding SHG imaging Z projections (scale bars, 100 μm). **(F)** Areas of collagen fibers in Control MO *versus cftr* MO regenerated fin as a function of tissue areas (left), and areas of collagen fibers in the regenerated fin of CF fishes following DMSO or curcumin exposure as a function of tissue areas (right). **(G)** Graph showing the ratio of area devoid of SHG fibers (from fiber ends to wound edge) following fin amputation. Ratios were determined by measuring the areas devoid of SHG fibers normalize with total regenerated tissue from F. **(H)** CT-FIRE-generated projections of SHG imaging of collagen fibers in the tail fin showing changes in organization of collagen fibers during tissue repair process. **(I-L)** Graphs showing quantitation of fiber number **(I)**, length **(J)**, width **(K)** as determined using CT-FIRE fiber analysis software (3 independent experiments, Two-way ANOVA with Tukey’s multiple comparisons test). **(L)** mRNA levels of *tgf-β* at 3dpA (relative gene expression from at least 3 independent experiments performed in triplicates, Two-way ANOVA with Tukey’s multiple comparisons test). **(M)** Graphs showing quantitation of fiber angle as determined using CT-FIRE fiber analysis software (3 independent experiments, Two-way ANOVA with Tukey’s multiple comparisons test). **(N)** 3D reconstruction from fin in cross section view revealing the spatial organization between tissue and the fibers (scale bars, 100 μm), with fibers found within the space between the 2 layers.

Together these data indicate that curcumin efficiently improves tissue repair and prevents tissue damage in CF animals, by restoring remodeling of collagen and tissue.

### Curcumin reduces inflammation and epithelial damage in CF patient-derived airway organoids

Finally, to further confirm the efficacy of curcumin in both reducing inflammation and tissue damage in CF, while extending our results into a human system, we validated the effect of curcumin on bronchiolar airway organoids (AOs)-derived from people with and without CF (10, 39). While CF-driven dysfunctional NRF2 activity was confirmed by the reduction in expression of *NRF2* in CF-AOs compared to healthy-AOs (10), we first showed that treatment with curcumin significantly increased *NRF2* expression in CF-AOs (**Figure 7A**). We showed that curcumin treatment brought *IL1β* and *IL8* expression back to normal level in CF-AOs (**Figure 7B**). Moreover, we evaluated the expression of genes related to the production and detoxification of ROS and observed that CF-AOs treated with curcumin exhibited a mitigated oxidative response associated with reduced expression of *DUOX1* and higher level expressions of *NQO1* (**Figure 7C**).

**Figure 7.**
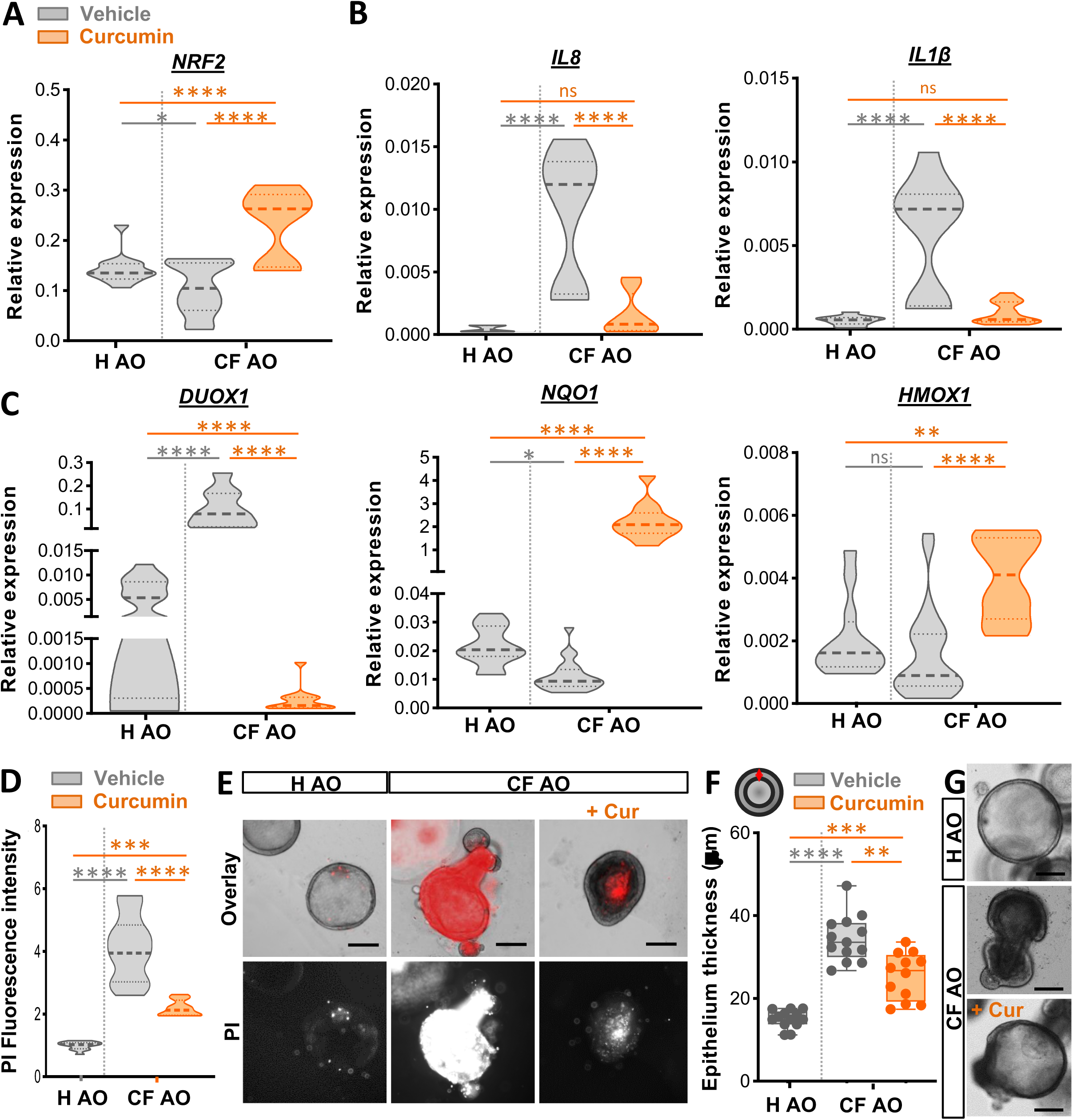
Treatment with curcumin alleviates inflammatory state and tissue damage in human CF airway organoid. (**A-C**) Healthy or CF airway organoid (AO) were incubated with vehicle (DMSO) control or 30 μM curcumin for 1 day. Gene expression levels of *NRF2* **(A)**, the pro-inflammatory cytokines *IL8* and *IL1β* **(B)**, the oxidative stress-related gene *DUOX1* and the *NRF2-*activated genes NQO1 and HMOX1 **(D)** were determined by RT qPCR (relative gene expression from at least 3 independent experiments performed in triplicates, ANOVA with Tukey’s multiple comparisons test). **(D-E)** Cell death analysis in AO treated with vehicle (DMSO) control or 30 μM curcumin for 4 days. Mean fluorescence intensity quantification of the propidium iodide (PI) incorporation **(D)** and representative images (scale bars, 300 μm) **(E)** in healthy and CF organoids treated or not with curcumin (n=12-13, from 3 independent experiments, Two-way ANOVA with Tukey’s multiple comparisons test). Data for at least three independent experiments are expressed as violin plots. **(F-G)** Quantification of the epithelial thickness (n=12-13, from 3 independent experiments, ANOVA with Tukey’s multiple comparisons test) **(F)** and representative images (scale bars, 300 μm) **(G)** in healthy and CF organoids treated or not with 30 µM of curcumin for 4 days.

As previously described, CF AOs display increased cell death and thicker epithelium compared to healthy ones (10), reflecting abnormal epithelial damage and remodeling. Having shown that curcumin efficiently limits tissue damage in CF zebrafish, curcumin treatment was next evaluated for its effects on cell integrity and mortality in AOs. Our results indicate that CF-AOs treated with curcumin displayed a significant decrease in cell mortality, when assessed by propidium iodide (PI) staining (**Figures 7D-E**). Furthermore, as shown in **Figures 7F-G**, treatment reduces wall thickness in CF AOs, denoting an improvement of cell integrity.

Altogether, these data support the potential of curcumin as a therapeutic for inflammatory lung damage in CF.

## Discussion

Over the last decade, research progress has led to spectacular advances in the management of CF, in particular with CFTR correctors and potentiators correcting the basic defect of CFTR function (40). However, excessive inflammation continues to be a major cause of lung destruction in CF, indicating a continued need for novel strategies to prevent inflammation-related pulmonary decline.

Among the defective mechanisms specific to CF-related inflammatory pathogenesis, the NRF2 signaling pathway, a master regulator of oxidative, inflammatory and regenerative responses, represent a promising target. Indeed, reports have shown differential expression of NRF2 expression and activation in several *in vitro* and *ex vivo* models of CF. Pharmacological activation of NRF2 results in decreased markers of oxidative and pro-inflammatory responses (9–12), suggesting that rebalancing NRF2 activity might simultaneously prevent inflammation and tissue damage in individuals with CF.

In the present study, using a combination of innovative animal and human models of CF, we sought to *i)* further explore the role of NRF2 in CF-related inflammatory status and associated tissue damage, and *ii)* determine the beneficial effect of curcumin, a known activator of NRF2, on inflammation and tissue repair in the context of CF. Our work identifies, for a first time, a role for NRF2 in neutrophilic inflammation, tissue damage and remodeling in CF. Importantly, we show that curcumin can restore normal level of inflammation and tissue integrity in a context of CF, by rescuing NRF2 signaling.

The link between NRF2 dysfunction and CF-associated neutrophilic inflammation remained elusive, although evidence suggests that impaired NRF2 activity correlates with excessive generation of ROS and pro-inflammatory cytokines in CF airway. Our results reveal that loss of Nfr2 signaling leads to an exuberant neutrophil response after injury in zebrafish. It coincides with an excessive production of epithelial ROS and pro-inflammatory cytokines IL8 and IL1β, all well-known key inducers of neutrophil mobilization (**Figure 2**). These data provide evidence that NRF2 pathway is instrumental to efficiently orchestrate neutrophil trafficking, likely by regulating oxidative and pro-inflammatory responses. Importantly, corroborating data from other CF models (9–12), Cftr-depleted zebrafish model also exhibits impaired Nrf2 signaling. Our findings confirm that impaired Nrf2 contribute to the neutrophilic inflammatory status in CF, since pharmacological NRF2 activation can block this effect in CF animals (**Figure 2**). The plausible explanation is that NRF2 dysfunction in CF leads to excessive production of ROS and pro-inflammatory cytokines by epithelia in response to injury or infection (or inherent to CFTR mutations), which in turn, cause exuberant neutrophilic inflammation.

Consistent with previous studies suggesting a role for NRF2 in tissue repair (26), we show that zebrafish lacking Nrf2 signaling exhibit incomplete tail fin regrowth after amputation, associated with disrupted collagen fiber formation and cell damage (**Figure 3**). Ablation of neutrophil in *nrf2*a-depleted animals partially rescues tissue repair, suggesting that neutrophilic inflammation is not the only cause of defective tissue repair in this context, and thus indicates that additional NRF2-mediated mechanisms, beyond its anti-inflammatory/antioxidant actions, are likely to participate in tissue repair processes. Similarly, we previously reported that Cftr-depleted zebrafish had abnormal tissue repair, partly due to an overactive inflammation (7). This also appears to be NRF2 dependent, since pharmacological activation of NRF2 signalling improves tissue repair in CF fish (**Figures 3,5-6**). These findings strongly suggest that deleterious changes in NRF2 activity are involved in defective tissue repair and tissue damage in CF.

Overall, these findings demonstrate the role of Nrf2 in CF-related inflammation and tissue damage, and thus provide a rationale for targeting this pathway to prevent inflammatory damage in patients. Curcumin, primary constituent in turmeric derived from *Curcuma Longa* rhizomes, is a potent activator of the NRF2 signaling pathway with anti-inflammatory, antioxidant and anti-infective therapeutic properties (41). We proposed here to study efficiency of curcumin on inflammatory response and tissue integrity in a context of CF, attempting to alleviate inflammation and tissue damage in CF.

We confirm that NRF2 activity is restored by curcumin treatment in both Cftr-depleted zebrafish and CF AOs (**Figures 4 and 7**). NRF2 is activated by stress (42); KEAP1 allows NRF2 to escape degradation, accumulate within the cell, and translocate to the nucleus, where it can bind to AREs promoting antioxidant and anti-inflammatory transcription programs. Interestingly, despite the large amounts of ROS produced in CF airways, NRF2 signalling fails to be activated and translocated to the nucleus and is rapidly degraded in CF (9, 11). Notably, several mechanisms could be proposed to explain the stimulating action of curcumin on NRF2 in a context of CF (15): by inhibiting KEAP1, by affecting the upstream mediators of NRF2 influencing the expression of *NRF2* and target genes, and/or by improving the nuclear translocation of NRF2.

Our results indicate that curcumin can significantly reduce CF inflammation *in vivo* and *ex vivo* by rebalancing excessive generation or inefficient detoxification of ROS, as well as elevated expression of *IL8* and *IL1β*. Consequently, curcumin alleviates CF-mediated neutrophilia. While curcumin has multiple effects and modes of action (41), we show that ablation of NRF2 signaling in CF zebrafish abolishes the protective effect of curcumin on CF-induced inflammation. Although we cannot exclude the possibility that curcumin modulates additional mechanisms to reduce inflammation, our conclusions provide evidence that curcumin prevents CF-mediated inflammation *via* activating the NRF2 pathway.

Since curcumin is safe and well-tolerated (43), these findings could have significant therapeutic implications for potently targeting inflammation in CF lung disease, and thus may serve as supplements to current therapeutic strategies or be an alternative to existing anti-inflammatory approaches. These data also suggest that curcumin may have beneficial effects on the extrapulmonary co-morbidities occurring in people with CF, such as gastro-intestinal and colorectal cancers (44, 45) or pancreas destruction and diabetes (46, 47). While CF is principally characterized by pulmonary disease, CF-related diabetes resulting from pancreatic islets destruction involving inflammation is a common feature among the individuals with CF and is associated with accelerated lung decline and increased mortality. CF pancreas is indeed subject to high level of ROS and IL1β, both known to cause β-cell apoptosis and implicated in both Type-1 and Type-2 diabetes. Furthermore, while pancreas destruction causes insulin insufficiency, chronic hyperglycemia is known to negatively affect pulmonary function, by impeding bacterial clearance, increasing oxidative stress and level of pro-inflammatory cytokines, and generating fibrosis in the lungs (48, 49). Importantly, an insufficient Nrf2 activity is often associated with the etiology of diabetes, and has been involved in the development of oxidative stress and inflammatory state in diabetic people (50). Interestingly, NRF2 activation protects mouse model of diabetes from ROS-induced damage (51), suggesting that targeting NRF2-modulated ROS production might show promise in the treatment of pancreas destruction in a context of CF. Thus, having shown promising results in the treatment of diabetes (52), by restoring normal level of inflammation, curcumin might also prevent diabetes in CF.

In CF, acute inflammatory responses disrupt airways integrity leading to abnormal tissue remodeling, increased accumulation and deposition of ECM components which can give rise to fibrosis (53). Our findings indicate that curcumin improves tissue repair in CF zebrafish, by reducing tissue damage and fibrotic phenotypes (**Figure 5-6**). The amelioration of tissue integrity is also observed in CF AO, associated with reduction of cell mortality and aberrant tissue remodeling (**Figure 7**), demonstrating that curcumin might prevent lung damage in people with CF. We have not yet identified the molecular events downstream of curcumin treatment in tissue repair and remodeling and how these might regulate collagen deposition. Modulation of macrophage (54) and fibroblast activity (55), two important cell populations in tissue repair processes including collagen deposition, could be plausible cellular targets.

Increasing evidence, including our previous works, demonstrates that hyper-susceptibility to infections in CF (56, 57) would be in part linked to defective bactericidal activities in professional phagocytes, themselves directly caused by CFTR dysfunction (58–60). Interestingly, while NRF2 is mostly known to be a regulator of oxidative and inflammatory responses, several studies indicate that NRF2 signaling pathway also plays a critical role in immune defense against pathogens (61). A defect in NRF2 in mice increases susceptibility to several CF-related bacteria, including *P. aeruginosa* (61, 62), *S. aureus* (63) and the mycobacterial species *Mycobacterium avium* (64), suggesting this could account for the infection phenotype in CF. Further investigations are needed to determine how CFTR/NRF2 interactions regulate antibacterial defense and how this defective axis contributes to increased susceptibility to infection in CF.

Anti-inflammatory therapies are of particular interest for CF lung disease but must be carefully studied to minimize the risk of impeding host immunity and thus worsening infection, as we recently observed with roscovitine (65). We showed that sulforaphane, another activator of NRF2, increases *M. abscessus* killing in several human models including CF AOs (10, 66), suggesting that activation of NRF2 might be a useful pharmacological approach for CF-associated defects in bacterial clearance. Hypothesizing that a decrease in Nrf2 signaling in people with CF indeed hampers their ability to defend against pathogens, it could be interesting to test whether activation of NRF2 by curcumin restores antimicrobial defense in CF. Moreover, while curcumin inhibits virulence factors in *P. aeruginosa*, such as the formation of biofilm or pyocyanin biosynthesis, gene involved in quorum sensing (67), the antibiotic synergic action of this compound also improves the activity of antibacterial agents against *S. aureus* (68), *M. abscessus* (69) and *P. aeruginosa* (70). Therefore, these results may have significant therapeutic implications for potently targeting inflammation in CF and improving host immunity to infection.

Because the low bioavailability of curcumin, associated with its poor solubility and absorption in free form in the gastrointestinal tract and its rapid biotransformation to inactive metabolites, there is a debate on its effectiveness and utility as a health-promoting compound (41). However, this study, involving inflammation and tissue damage management in CF, was conducted with a free molecule, thus supporting its therapeutic and protective effects. With the availability of highly bioavailable curcumin formulations (71), it should be now possible to exploit the full activities and benefits of curcumin in CF, and clinical trials will be necessary to confirm its effects in humans.

To conclude, our findings indicate that rescuing NRF2 using curcumin may be a targeted therapeutic strategy to both mitigate inflammation and restore tissue repair, and thus prevent inflammatory damage in individuals with CF.

## Acknowledgments

We acknowledge the INRAE Infectiology of Fishes and Rodents Facility (IERP-UE907, Jouy-en-Josas Research Center, France doi.org/10.15454/1.5572427140471238E12) and the ZEFIX (ZEbraFIsh and Xenopus platform, Lphi, University of Montpellier) fish facilities, and Dimitri Rigaudeau, Penelope Simon, Catherine Gonzalez and Victor Goulian for zebrafish maintenance and care, with a special thanks to Magalie Bouvet for technical assistance. IERP Facility belongs to the National Distributed Research Infrastructure for the Control of Animal and Zoonotic Emerging Infectious Diseases through In Vivo Investigation (EMERG’IN DOI: doi.org/10.15454/90CK-Y371). We thank the IERP for giving access to their injection platforms, the zebrafish phenotyping platform of IERP and imaging facility of MRI (Montpellier Ressources Imagerie), and Elodie Jublanc and Vicki Diakou for their assistance. We also thank Philippe Clair, the manager of the quantitative PCR facility of the University of Montpellier. We also thank the University of Sheffield, the University of Versailles Saint-Quentin, the University of Montpellier, the animal health division of INRAE and the University of Toulouse for support.

## Notes

**Acknowledgments**: This study was supported by the Horizon 2020 Research and Marie Skłodowska-Curie Innovation Framework Program (H2020-MSCA-IF-2016, CFZEBRA (751977); AB), the Fondation pour la Recherche Médicale (ARF201909009156; AB), the FC3R through the ZFishforCFCare project (22FC3R-017; AB), the CF Trust (Workshop funding (160161); Strategic Research Centre SRC018; AB, RAF and SAR), the Vaincre La Mucoviscidose and Grégory Lemarchal foundations (RF20210502852/1/1/48; CC and SALI). LY is supported by the Horizon 2020 Program Marie Skłodowska-Curie Innovative Training Network (H2020-MSCA-ITN-2020, INFLANET (955576)). SC was the recipient of a PhD fellowship from the French Ministry of Research.

### Competing Interest Statement

The authors have declared no competing interest.

