## Supplemental File for "Curcumin-mediated NRF2 induction limits inflammatory damage in preclinical models of cystic fibrosis"

### Supplementary Data

#### Supplementary figures

**Figure S1. Analysis of the protein sequence conservation between human NRF2 and the two zebrafish orthologs, Nrf2a and Nrf2b.** Protein sequences of human NRF2 (hNRF2) (Uniprot Q16236), zebrafish Nrf2a (Uniprot Q7ZVI2) and zebrafish Nrf2b (Uniprot G8DKA8) were aligned using Clustal Omega from the EMBL-EBI server (1). Analysis of the sequence conservation was done with ALINE (2). Residue numbering corresponds to the sequence of hNRF2. Secondary structure elements derived from the AlphaFold (3) structural model predicted for hNRF2 are indicated above the sequence, α-helices are shown in light blue, β-strands in light pink. The position of the various Neh domains in hNRF2 protein sequence are indicated as colored boxes above the sequences, using the same domain color code as in main Figure 1.

**Figure S2. Structural analysis of zebrafish Nrf2b and comparison with human NRF2. (A)** Predicted domain organization of zebrafish Nrf2b protein. Six Neh domains are conserved with hNRF2 and zebrafish Nrf2a, only Neh4 is absent. **(B)** 3D-structure of zebrafish Nrf2b predicted with AlphaFold. As for NRF2 and Nrf2a, Nrf2b is proposed to adopt a 3D-fold bearing long unstructured regions that alternate with shorter regions displaying secondary structure elements, mostly α-helices. The Neh domains are represented with the same color code as in panel **A**. **(C)** Structural overlay of the AlphaFold-predicted model for zebrafish Nrf2b (dark purple) with that of hNRF2 (cyan). The position of the secondary structure elements within the Nrf2b model seems to diverge more from hNRF2 than in the case of Nrf2a. **(D)** Structural overlay of the Neh1 domains from AlphaFold-predicted Nrf2b structure (dark purple), from AlphaFold-predicted hNRF2 structure (cyan), and from the X-ray structure of the hNRF2-MafG complex (salmon; PDB ID 7X5F; (4)). **(E)** Structural overlay of the Neh2 domains from AlphaFold-predicted Nrf2b structure (dark purple), from AlphaFold-predicted hNRF2 structure (cyan), and from the X-ray structure of the human NRF2-Keap1 complex (salmon; PDB ID 3WN7; (5)).

**Figure S3. Basal neutrophil number in *nrf2a*- and *nrf2b*-depleted zebrafish larvae compared to control animals. (A)** Total number of neutrophils in whole control morphant (control MO), *nrf2a* morphant (*nrf2a* MO) and *nrf2b* morphant (*nrf2b* MO) *TgBAC(mpx:EGFP)i114* (6) at 3 days post-fertilization (dpf). Each dot represents the total number of neutrophils in a single larva (from 2 independent experiments; One-way ANOVA with Dunnett's post-test). **(B)** Representative photomicrographs of neutrophilic distribution and number in control MO, *nrf2a* MO and *nrf2b* MO

*TgBAC(mpx:EGFP)i114* larvae following tail fin amputation (scale bars, 200 μm). Nrf2a ablation has no measurable impact on the total number of neutrophils, indicating that elevation in neutrophil numbers observed at the site of injury is not due to an overall increase in neutrophils within *nrf2a*-depleted zebrafish.

**Figure S4. Loss of Nrf2a promotes exuberant oxidative and pro-inflammatory responses in zebrafish model. (A)** Control MO and *nrf2a* MO *TgBAC(mpx:EGFP)i114* were pretreated with DPI or DMSO (as control) prior tail fin amputation procedure, then injured and immediately put back in treatments. The number of neutrophils at wound was counted at 2 hours post-amputation (hpA) (3 independent experiments, Two-way ANOVA, Bonferroni's multiple comparisons test). **(B-F)** Caudal fin amputations (dotted lined) were performed distally without injury to the notochord at 3 dpf. **(C-D)** Control MO and *nrf2a* MO were stained with CellROX® to label ROS production. Larvae were then caudal fin amputated and oxidative activity observed and analyzed at 30 min post-amputation (mpA) under a fluorescent microscope. Relative ROS intensity as violin plots (Relative fluorescence intensity (RFI), n=18 from 3 independent experiments; Mann-Whitney U test) **(C)** and associated pseudocolored photomicrographs of injured fins **(D)** revealing oxidative activity at the wound-edge (50 μM from the amputation edge (AE) (dotted line); scale bar, 100 μm). **(E-F)** Control MO and *nrf2a* MO were generated in *Tg(il1b:eGFP-F)ump3* line*,* allowing to visualize *Il1β* expression (7). Larvae are caudal fin amputated then the expression of *Il1β* was observed and analyzed at 2 hpA under a fluorescent microscope. Relative *Il1β* intensity as violin plots (n=18 from 3 independent experiments; Two-tailed Student *t*-test) **(E)** and associated pseudocolored photomicrographs of injured tails (50 μM from the AE; scale bar, 100 μm) **(F)** revealing *Il1β* expression at the wound-edge. **(G)** Control MO and *il1β* morphant (*il1β* MO, generated using the validated *il1β* morpholino (7)) *TgBAC(mpx:EGFP)i114* were tail fin amputated then the number of neutrophils at wound was counted at 4 hpA (2 independent experiments, Two-tailed Student *t*-test). Deletion of Il1β signaling reduces neutrophilic response in zebrafish.

**Figure S5. *in vivo* tissue repair performance assessment in the absence of Nrf2b signaling.** 2 dpf control MO and *nrf2b* MO were tail fin amputated, then the potential of tissue repair was evaluated by measuring regenerated tissue at 3 days post-amputation (dpA). Measurement of regenerated fin areas (left), length (middle) and width (right) (n=18 from 3 independent experiments, Two-tailed Student *t*-test). Error bars represent standard error of the mean (SEM).

**Figure S6. Influence of neutrophilic inflammation on tissue repair performance of Nrf2a-depleted zebrafish.** Ablation of neutrophil was carried out in control MO and nrf2a MO by injecting the *csf3r* morpholino (8). Embryos were then tail fin amputated at 2 dpf, and the potential of tissue repair was evaluated by measuring regenerated fin areas at 3 dpA in the presence or absence of neutrophils (n=14-16 from 2 independent experiments, Two-way ANOVA, Tukey's multiple comparisons test). Error bars represent SEM. Removal of neutrophils slightly restores fin regrowth in *nrf2a* MO, suggesting that exuberant neutrophilic inflammation is partially involved in impaired tissue repair associated with the loss of Nrf2a.

**Figure S7. Curcumin restores Nrf2b signaling in *cftr*-depleted zebrafish.** Control MO and *cftr* MO were pre-treated with DMSO or curcumin prior to caudal fin amputation procedure, then injured and immediately put back in treatments until analysis. mRNA levels of *nrf2b* at 2 hpA (relative gene expression from at least 3 independent experiments performed in triplicates, Two-way ANOVA with Tukey's multiple comparisons test).

**Figure S8. Curcumin does not influence the resolution phase of neutrophilic inflammation.** In order to assess the pro-resolution activity of curcumin, control MO and *cftr* MO *TgBAC(mpx:EGFP)i114* were tail fin amputated and treated from 4 hpA with 2.5 μg/mL of curcumin or DMSO as control vehicle. Neutrophil number at the wound was counted at 8 hpA (n=14-18 from 2 independent experiments, Two-way ANOVA with Tukey's multiple comparisons test). Similar wound-associated neutrophil numbers were observed in curcumin- and DMSO-treated larvae, suggesting that persistence of neutrophil inflammation was not affected by curcumin exposure.

**Figure S9. Curcumin exposure alleviates neutrophilic inflammation in CF zebrafish in a context of bacterial infections. (A-B)** *cftr* MO *TgBAC(mpx:EGFP)i114* zebrafish were injured and infected with fluorescent *Staphylococcus aureus* (Sa), *Pseudomonas aeruginosa* (Pae) or *Mycobacterium abscessus* (Mabs). **(B)** Neutrophil number at the infected wound was counted at 2 hpA (3 independent experiments, One-way ANOVA with Dunnet's multiple comparisons test). **(C)** *cftr* MO were pre-treated with DMSO or curcumin prior to caudal fin amputation procedure, then injured, and immediately exposed with *S. aureus*, *P. aeruginosa* or *M. abscessus* for 1 hour. Double injured/infected larvae were then put back in treatments until analysis. Neutrophil number at the infected wound was counted at 2 hpA (3 independent experiments, Two-tailed Student *t*-test).

**Figure S10. Action of curcumin in absence of Nrf2 signaling.** Control MO, nrf2a MO, nrf2b MO and nff2a+b MO were pre-treated with DMSO or curcumin prior to caudal fin amputation procedure, then injured and immediately put back in treatments until analysis. Neutrophil number at the wound at 2 hpA (3 independent experiments, Two-way ANOVA with Bonferroni's multiple comparisons test) in the absence or presence of curcumin.

**Figure S11. Exuberant wound-induced Il1β signaling drives the overactive neutrophil response in CF zebrafish. (A-B)** Il1β activity upon wounding in control and *cftr* MO *Tg(il1b:eGFP-F)ump3* (7). Control and *cftr* MO *Tg(il1b:eGFP-F)ump3* were caudal fin amputated then the activity of *Il1β* was observed and analyzed at 2 hpA. Relative *Il1β* intensity as violin plots (n=18 from 2 independent experiments; Mann-Whitney U test) **(A)** and associated pseudocolored photomicrographs of injured tails **(B)** revealing *il1β* signaling at the wound-edge (scale bar, 100 µm). Loss of CFTR leads to an increase in *il1β* activation in response to injury. **(C)** Inhibition of *Il1β* was carried out in CF animals by injecting the *Il1β* morpholino (7). Neutrophil number at the wound at 2 hpA (3 independent experiments, Two-way ANOVA with Tukey's multiple comparisons test).

**Figure S12. Curcumin exposure reduces tissue damage and improves tissue repair in CF zebrafish in a context of bacterial infection. (A-B)** 2dpf *cftr* MO *TgBAC(mpx:EGFP)i114* were injured and infected with fluorescent *S. aureus*, *P. aeruginosa* or *M. abscessus* for 4 h. The potential of tissue repair was evaluated by measuring regenerated fin areas at 3 dpA (n=21 from 3 independent experiments, One-way ANOVA with Dunnet's multiple comparisons test). **(C)** *cftr* MO were pre-treated with DMSO or curcumin prior to caudal fin amputation procedure, then injured, and immediately exposed with *S. aureus*, *P. aeruginosa* or *M. abscessus* for 1 hour. Double injured/infected larvae were then put back in treatments until regenerated fin areas analysis at 3 dpA (n=21-24 from 3 independent experiments, Two-tailed Student *t*-test).

**Figure S13. Curcumin restores collagen fiber and mesenchymal cell distribution in Cftr-depleted animals. (A)** Second harmonic generation (SHG) and high magnification bright field imaging showing collagen fibers signature in control MO *versus cftr* MO. Z-projections of caudal fins imaged at 3 dpA (scale bar, 100 µm). **(B)** control MO and *cftr* MO were pre-treated with DMSO or curcumin prior to caudal fin amputation procedure, then injured and immediately put back in treatments for 4 h. Z-projections of caudal fins revealing collagen fiber organization at 3 dpA (scale bar, 100 µm). **(C)** Control MO and *cftr* MO are generated in the *Tg(rcn3:GAL4/UAS:mCherry)* line labelling mesenchymal cells (9). Confocal fluorescence of maximum intensity projection of unamputated tail fins showing mesenchymal cell distribution and morphology (top panel, scale bar, 100 µm). *Tg(rcn3:GAL4/UAS:mCherry)* control MO and *cftr* MO were pre-treated with DMSO or curcumin prior to caudal fin amputation procedure, then injured and immediately put back in treatments for 4 h.

Mesenchymal cell distribution and morphology was observed at 3 dpA in confocal microscope (scale bar, 100 µm).

#### Tables

**Table 1. Minimum Inhibitory Concentrations of Curcumin against *M. abscessus*, *P. aeruginosa* ans S. *aureus***

| Bacteria | Cation-adjusted Muller-Hinton Broth (μM) |
|---|---|
| *M. abscessus* | > 256 |
| *P. aeruginosa* | > 256 |
| *S. aureus* | > 256 |

**Table 2. RT-qPCR primers used in the study**

| **Gene** (NCBI RefSeqGene) | **Primers 5'-3'** | **Reference** |
|---|---|---|
| | **Zebrafish model** | |
| ***ef1a*** (NM_ 131263) | fw TCTGTTACCTGGCAAAGGG<br>rev TTCAGTTTGTCCAACACCCA | |
| ***nrf2a*** (NM_182889) | fw GAGCGGGAGAAATCACACAGAATG<br>rev CAGGAGCTGCATGCACTCATCG | (10) |
| ***nrf2a*** (NM_001257183) | fw GGCAGAGGGAGGAGGAGACCAT<br>rev AAACAGCAGGGCAGACAACAAGG | (10) |
| ***duox*** (XM_021470722) | rev GTTGGCTTTGGTGTAACTGTA<br>fw GCCCAGGCTGTGAGAG | (11) |
| ***cxcl8*** (XM_001342570) | fw CCTGGCATTTCTGACCATCAT<br>rev GATCTCCTGTCCAGTTGTCAT | (12) |
| ***Il1β*** (NM_212844) | fw TGGACTTCGCAGCACAAAATG<br>rev GTTCACTTCACGCTCTTGGATG | (12) |
| ***hmox1a*** (NM_001127516) | fw TAAAAACGAAGTGGGGCGGT<br>rev TGTTCAGACAGATCACTGCCA | (13) |
| ***sod2*** (NM_199976) | fw ATGCTGTGCAGAGTCGGATATG<br>rev GCTGAAGGGAGACTTGGGTT | (14) |
| ***nqo1*** (NM_001204272) | fw CTTGATCGCAGAAGAATAATATGC<br>rev CAGCACTCCATTCTGTAAGGG | (15) |
| | **Human organoid model** | |
| ***GAPDH*** (NM_002046) | fw CTCCAAAATCAAGTGGGGCGATG<br>rev GGCATTGCTGATGATCTTGAGGC | (16) |
| ***NRF2*** (NM_006164.5) | fw TCAGCGACGGAAAGAGTATGA<br>rev CCACTGGTTTCTGACTGGATGT | PrimerBank |
| ***DUOX1*** (NM_017434.5) | fw TTCACGCAGCTCTGTGTCAA<br>rev AGGGACAGATCATATCCTGGCT | (17) |
| ***DUOX2*** (NM_014080.5) | fw ACGCAGCTCTGTGTCAAAGGT<br>rev TGATGAACGAGACTCGACAGC | (17) |

| | | |
|---|---|---|
| ***IL8*** (NM_000584) | fw TACTCCAAACCTTTCCACCCC<br>rev CTTCTCCACAACCCTCTGCA | (16) |
| ***IL1β*** (NM_000576) | fw AGCTACGAATCTCCGACCAC<br>rev GGGAAAGAAGGTGCTCAGGTC | (16) |
| ***HMOX1*** (NM_002133.3) | fw TCCGATGGGTCCTTACACTC<br>rev TAAGGAAGCCAGCCAAGAGA | (18) |
| ***SOD2*** (NM_000636.4) | fw TTTCAATAAGGAACGGGGACAC<br>rev GTGCTCCCACACATCAATCC | PrimerBank |
| ***NQO1*** (NM_000903.3) | fw CAGACGCCCGAATTCAAATC<br>rev AGGCTGCTTGGAGCAAAATACA | (18) |

**Table 3. Information of the consented donors**

| **Donor** | **Sex** | **Age** (years) | **Pathology** | **Mutation** |
|---|---|---|---|---|
| Healthy 1 | Female | 62 | Lung cancer | - |
| Healthy 2 | Female | 64 | Lung cancer | - |
| CF 1 | Female | 30 | Cystic fibrosis | G542X -1811+1.6kb A→G |
| CF 2 | Male | 34 | Cystic fibrosis | ΔF508 / N1303K |

#### Methods

##### Zebrafish Husbandry and Ethics Statement

Zebrafish (*Danio rerio*) experiments described in the present study were performed by authorized staff and conducted by following the 3Rs -Replacement, Reduction and Refinement- principles in compliance with the European Union guidelines for handling of laboratory animals. Breeding and maintenance of adult fish were performed in the Bateson Centre (University of Sheffield, Sheffield, UK; license number P1A4A7A5E), the IERP (Inrae, Jouy-en-Josas, France; license number C78-720) and the ZEFIX-Lphi (CNRS, University of Montpellier, Montpellier, France; license number CEEA-LR-B34-172-37) fish facilities, according to the local animal welfare standards set approved by the UK Home Office under Animal Welfare and Ethical Review Body, the Directions Sanitaires et Vétérinaires de Versailles et de l'Hérault et Comités d'Ethiques pour l'Expérimentation Animale de Paris-Saclay et de la région Languedoc Roussillon (France) and the French Ministry of Agriculture and Food.

Experimental procedures were performed using the pigment-less *nacre* (19) or golden (20) lines along with the following transgenic lines : *TgBAC(mpx:eGFP)i114* (6) to label neutrophils; *Tg(il1β:eGFP-F)ump3* (7) and *Tg(pNF-κB:EGFP)sh235* (21) to visualize the transcriptomic expression of *il1β and nf-κb* reciprocally; *Tg(rcn3:gal4)$^{pd1023}$* and *Tg5(UAS:mCherry)$^{pd1112}$* referred to as *Tg(rcn3:gal4/UAS:mCherry)* to label mesenchymal cells (9).

All zebrafish experiments were performed on larvae <5 days post-fertilization (dpf). The number of animals used for each procedure was guided by pilot experiments or by past results (22, 23). Zebrafish eggs were obtained from pairs of adult fish by natural spawning and raised in E3 water (24) at 28°C and exposed on a 14:10 hours light/dark cycle to maintain proper circadian conditions. For zebrafish anesthesia procedures, larvae are immersed in a 168 mg/L tricaine (Sigma-Aldrich) or 0.0075 % eugenol (Fisher Scientific) solution in E3 water. When required, larvae were cryo-anesthetized by incubation on ice for 10 minutes and then euthanized using an overdose of tricaine (500 mg/L).

##### Morpholino Injections

Morpholinos used in this study were purchased from Gene Tools. The morpholinos for *cftr* knock-down (5'-GACACATTTTGGACACTCACACCAA-3'), *nrf2a* knock-down (5'-CATTTCAATCTCCATCATGTCTCAG-3'), *nrf2b* knock-down (5'-AGCTGAAAGGTCGTCCATGTCTTCC-3') and *il1β* (5'-CCCACAAACTGCAAAATATCAGCTT-3') were prepared and injected into one-cell-stage zebrafish as described (7, 10, 25). For the selective depletion of neutrophils into zebrafish larvae, *csf3r* morpholino (5'-GAAGCACAAGCGAGACGGATGCCAT-3') targeting the *csf3* gene was used (8). A standard control morpholino (5'-CCTCTTACCTCAGTTACAATTTATA-3') was used as a negative control.

**Drug Treatments in Zebrafish larvae**

To explore the therapeutic efficacy of curcumin, zebrafish larvae were incubated in sterile E3 medium supplemented with 2.5 µg/mL of curcumin (Sigma-Aldrich). Both ML 385 (Sigma-Aldrich) and CDDO-Methyl ester (CDDO-Me) (Sigma-Aldrich) were used at 1.5 µg/mL working concentration to reciprocally inhibit or activate Nrf2 signaling. The doses we identified that did not generate toxicity. Oxidative activity was blocked using 100 µg/mL of diphenyleneiodonium (DPI, Sigma-Aldrich) as described earlier (26). Dimethyl sulfoxide (DMSO, Sigma-Aldrich) was used as vehicle control.

**RNA Isolation from Zebrafish larvae and qRT-PCR Analysis**

Total RNA from a pool of 10-15 larvae per biological experiment was extracted using Nucleospin RNAII kit (Macherey-Nagel) at time points indicated in Figure legends and cDNA synthesized with M-MLV reverse transcriptase (Invitrogen). Real-time RT-PCR were performed using SensiFAST SYBR Green No-ROX mix (Thermo Fisher Scientific) on a LightCycler 480 instrument (Roche) as described (27) and gene expressions were detected with the gene-specific primers provided in **Table 2**. Each experiment was run in triplicate. ΔCT was calculated using the housekeeping gene *ef1*α as a reference gene. Relative expression levels were calculated using the $2^{-\Delta Ct}$ method.

**Inflammation Assays in Zebrafish larvae**

Inflammation was elicited by distal or proximal tail fin amputation on 3 dpf anesthetized larvae using a microscalpel (5 mm depth; World Precision Instruments) according to established procedures (6, 22). Neutrophil response was observed and evaluated by manually assessing the number of cells at wound sites at relevant time points using a fluorescence dissecting stereomicroscope (Leica) as previously defined (22): 2 hours post-amputation (hpA) (acute phase of neutrophil response), 4 hpA (neutrophil recruitment peaking) or 8 hpA (resolution phase of neutrophil response). Neutrophils at the wound sites were imaged on an Eclipse TE2000 U inverted compound fluorescence microscope (Nikon UK Ltd., Kingston upon Thames, UK) equipped with a 20x NA objective lens, using a 1394 ORCA-ERA camera (Hamamatsu Photonics Inc). Overlays of fluorescent and DIC images were assembled using FIJI (National Institutes of Health).

Epithelial *il1*$\beta$ and *nf-*$\kappa$*b* expressions at the wound sites (50 µm anterior from the wound marge) were observed and captured at 2 hpA, using an Olympus MVX10 fluorescence microscope (Olympus, Life Science) equipped with a X-Cite Xylis LED (Excelitas Technologies) light source. Images were acquired with a Hamamatsu ORCA-spark Digital CMOS C11440-36U camera (Hamamatsu Photonics Inc) and processed using Olympus CellSens Standard 3.1 software (Olympus, Life Science). *il1*$\beta$ and *nf-*$\kappa$*b* activities were assessed in FIJI using the intensity of fluorescence and normalized to uninjured larvae (control or morphant animal; untreated or treated animal).

**Oxidative Activity Assay in Zebrafish larvae**

Epithelial oxidative response was detected using CellROX® deep red or CellROX® green (Thermo Fisher Scientific) following protocol previously established (22) and elicited by tail fin amputation as described above or by laser-mediated injury (23).

For tail fin amputation procedure, following CellROX® green staining, larvae were injured then reactive oxygen species (ROS) production at the wound sites (50 μm anterior from the wound marge) were observed and captured at 30 minutes post-amputation (mpA) using fluorescence microscope (Olympus, Life Science). For laser injury, following CellROX® deep red staining, anaesthetized larvae were mounted in 0,8% low melting point agarose (Thermo Fisher Scientific). A femtosecond titanium:sapphire pulsed laser (Coherent Vision II) set at 800-nm and at 40% laser power was used to injure caudal fin. A 9.23 µm x 9.23 µm field of view was scanned during 1 second with a pixel well time of 72 ns using an upright Leica SP8 confocal microscope equipped with an HCX IRAPO 25X/0.95NA water immersion objective (Leica Microsystems). Then ROS production at the wound sites was immediately time-lapsed at indicated time points throughout acute inflammatory stage.

The laser makes a small circular injury to tissue resulting in the production of ROS and accumulation of neutrophils in this area (23). CellROX® deep red was excited with a 633 nm laser, then fluorescence and bright field transmission images were acquired with a with photomultiplier tube detectors and processed using LASX software (Leica). Oxidative activity was assessed in FIJI using the intensity of fluorescence and normalized to uninjured larvae (control or morphant animal; untreated or treated animal).

***In vivo* tissue repair assay**

Tissue repair assay was performed following established methods (22). Briefly, 2 dpf embryos were anesthetized then tail fins amputated at the boundary of the notochord without injury to the notochord (distal injury). Tissue repair performances were evaluated by assessing the regenerated tail fin area/length/width and wound contour length at 3 days post-amputation (dpA) under a stereomicroscope (MZ10F, Leica Microsystems) equipped with a PLANAPO 1X objective lens. Image were acquired with a MC170HD camera (Leica), then processed and analyzed with FIJI: regrowth area was measured by outlining the total fin tissue area distal to the notochord using the polygon tool, regrowth length and width were measured by using the straight-line tool. Percentage of regeneration was calculated by normalizing the regenerated tail fin areas/length/width *versus* fin areas/length/width of unamputated animals (control or morphant animal). To assess the length of wound edge as an indicator of projections, the length of the wound edge was measured using the freehand line tool to trace the contour of the wound edge, as previously described (28).

**Bacterial strains, growth conditions and tail wound infection challenges**

*Staphylococcus aureus* (SH1000 strain carrying pMV158-mCherry (29), generously provided by Simon Foster (Florey Institute, Sheffield, UK)), *Pseudomonas aeruginos*a (PAO1 strain) and *Mycobacterium abscessus sensu stricto* (CIP104536$^T$ strain, morphotype smooth (S)), were used for this study.

*S. aureus* were grown using Brain Heart Infusion (BHI) media (MilliporeSigma) with tetracycline 5 µg/ml (Sigma-Aldrich). *P. aeruginosa* were grown using Luria-Bertani media (LB Miller, MilliporeSigma). *M. abscessus* were grown in Middlebrook 7H10 Agar and Middlebrook 7H9 Broth media (MilliporeSigma), supplemented with 10% acid/albumin/dextrose/catalase (ADC) enrichment (MilliporeSigma). To prepare *S. aureus* and *P. aeruginosa* inoculates, 1 ml of appropriate medium was inoculated with a fresh colony of bacteria and incubated at 37°C overnight with shaking. 500 µl of this overnight culture was then added to 10 ml of appropriate medium and incubated at 37°C with shaking to achieve growth to mid-logarithmic phase ($OD_{600}$ ≈ 0,6-0,8). *M. abscessus* inoculates were prepared as previously described (30). Next, to generate infected E3 medium, bacteria were harvested by centrifugation, washed with Phosphate-Buffered Saline (PSB, Gibco™, Thermo Fisher Scientific), and resuspended at an $OD_{600}$ of 1 in sterile E3 water. Caudal tail fin amputation of anesthetized-larvae was performed as described above, then injured larvae were immediately transferred in sterile or infected E3 medium for 1 hour at 28°C. Uninfected and infected larvae were then rinsed three times with sterile E3 medium to wash away bacteria, and maintained at 28°C until imaging for neutrophil mobilization or wound healing assays as described above.

**Minimum Inhibitory Concentration Determination**

Minimum Inhibitory Concentrations (MICs) were determined using the microdilution method, in cation-adjusted Mueller-Hinton broth, according to the Clinical and Laboratory Standards Institute (CLSI) guidelines (31). Serial 10-fold dilutions of log-phase cultures were plated and incubated at 37°C for 3 to 4 days, and the MICs were recorded by visual inspection and defined as the minimum concentration required to inhibit 99% of the growth. Experiments were done in triplicate in three independent occasions.

**Second Harmonic Generation Microscopy and Collagen Fiber Observation**

Second harmonic generation (SGH) imaging was performed on living zebrafish larvae mounted in 0.8% low-melting agarose (Sigma-Aldrich), using an upright Leica SP8 two-photon microscope equipped with an HCX IRAPO 25X/0.95NA water immersion objective (Leica Microsystems). Caudal fin areas were excited at 1040 nm with a Chameleon Vision II laser (Coherent). SHG signal, corresponding to collagen fibers, was detected in a non-descanned (NDD) pathway with a hybrid detector (HyD, Leica) associated with a 525/50 nm bandpass filter (green channel). Images of regenerated fins were acquired with photomultiplier tube detectors and processed using LASX software (Leica). Overlays of fluorescent and Brightfield images, and maximal projections of image stacks were assembled using FIJI. 3D reconstitution was produced using Imaris 9.0 software (Bitplane).

*Collagen fiber quantification.* Collagen alignment quantification was performed using the collagen quantification platform CurveAlign 4.0 (https://eliceirilab.org/software/curvealign/) (32).

*Area devoid of collagen fibers*. To measure the area between the wound tissue edge and the ends of the SHG detected fibers, maximum projections of SHG z-stacks, with the notochord vertically centered, were generated. A freehand line was drawn along the ends of the fibers and applied to the corresponding brightfield image, then the area between this fiber-end boundary and the wound edge was drawn and measured using FIJI software.

**Human Bronchiolar Airway Organoid Ethics Statement, Preparation and Treatment**

The acquisition of patient data and lung tissue for human bronchiolar airway organoids (AOs) generation was performed in accordance with the Medical Research Involving Human Subjects Act and approved by the CHU of Toulouse and the CNRS under the number agreements CHU/19244C and CNRS/205782. Healthy tissue from two volunteers with lung cancer and lung biopsies from two consented patients with CF (**Table 3**) were used to derive AOs as previously described (16, 33, 34).

To explore the therapeutic efficacy of curcumin, healthy-AOs and CF-AOs were embedded in 40 µl drops of Cultrex growth factor reduced basement membrane extracts (BME) type 2 (R & D Systems), and seeded on Nunclon Delta surface 24-well plates (Thermo Fisher Scientific). Once the Cultrex has polymerized, 500 µl of AO complete media, without N-acetylcysteine and without antibiotics, were added. According to the indicated conditions, AOs were stimulated or not with 30 µM curcumin for 1 or 4 days.

**Human Bronchiolar Airway Organoid Mortality Assay**

To determine AO mortality, healthy-AOs and CF-AOs stimulated or not for 4 days with 30 µM curcumin were stained with 50 µg/ml Propidium Iodide (PI, Thermo Fisher Scientific) as previously described (34). PI incorporation was then immediately analyzed by live imaging using an EVOS M7000 Imaging System (Thermo Fisher Scientific) (10x, at 37°C with 5% $CO_2$) and cellular death was assessed in FIJI using the intensity of fluorescence.

**Human Bronchiolar Airway Organoid Epithelial Thickness**

Healthy-AOs and CF-AOs were embedded in a Cultrex matrix, then seeded and cultivated for a minimum of five weeks on Nunclon Delta surface 24-well plates, utilizing complete airway organoid media. The organoids were maintained at 37°C with 5% $CO_2$. Upon completion of the cultivation period, the organoids were either treated with 30 µM of curcumin for 4 days or left untreated. Following this treatment, bright-field images of each well were acquired using an EVOS M7000 Imaging System (10x magnification), and subsequently, the analysis of epithelial thickness for each organoid was conducted using the FIJI software.

**RNA Isolation from Human Bronchiolar Airway Organoids and qRT-PCR Analysis**

Healthy- or CF-AO (~15 per condition) stimulated or not with 30 µM curcumin for 1 day, were harvested with cold 1X phosphate-buffered saline (PBS). The organoids collected were centrifugated at 800g x 5 minutes, and the supernatants were discarded. Total RNA was extracted using the RNeasy mini kit (Qiagen), followed by retrotranscription (150 ng) with the Verso cDNA Synthesis Kit (Thermo Fisher Scientific). mRNA expression was evaluated with an ABI 7,500 real-time PCR system (Applied Biosystems) and the SYBRTM Select Master Mix (Thermo Fisher Scientific). Relative quantifications were determined by the $2^{-\Delta Ct}$ method and normalized to GAPDH. Primer sequences are provided in **Table 2**.

**3D-modeling and Structural Analysis**

The 3D structural models for the zebrafish *nrf2a* and *nrf2b* were predicted using AlphaFold (v2.3.2) (3) from the online Colab notebook. Structural alignment was done in Coot (35) and all figures were made with the Pymol Molecular Graphics System (version 0.99rc6, DeLano Scientific LLC). Sequence alignment were done in Clustal Omega (1) and sequence conservation was analyzed with ALINE (2).

**Quantification and statistical analysis**

Statistical analysis was performed using Prism 10.0 (GraphPad Software) and detailed in each Figure legend. ns, not significant ($p \geq 0.05$); *$p<0.05$; **$p<0.01$; ***$p<0.001$; ****$p<0.0001$.

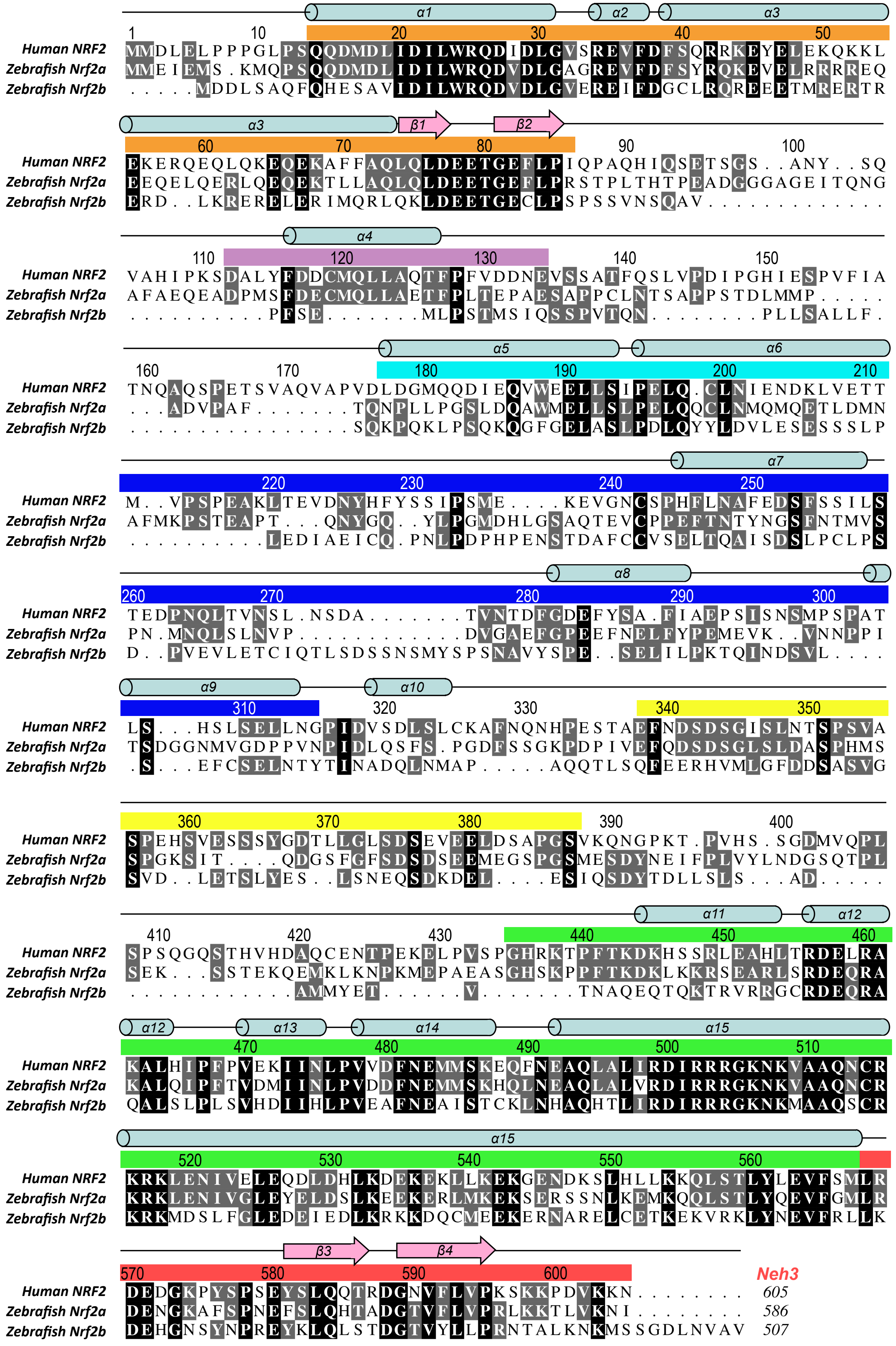
Neh2
Neh4
Neh5
Neh7
Neh6
Neh1
Neh3
α1
α2
α3
α4
α5
α6
α7
α8
α9
α10
α11
α12
α13
α14
α15
β1
β2
β3
β4
Human NRF2
Zebrafish Nrf2a
Zebrafish Nrf2b

S1

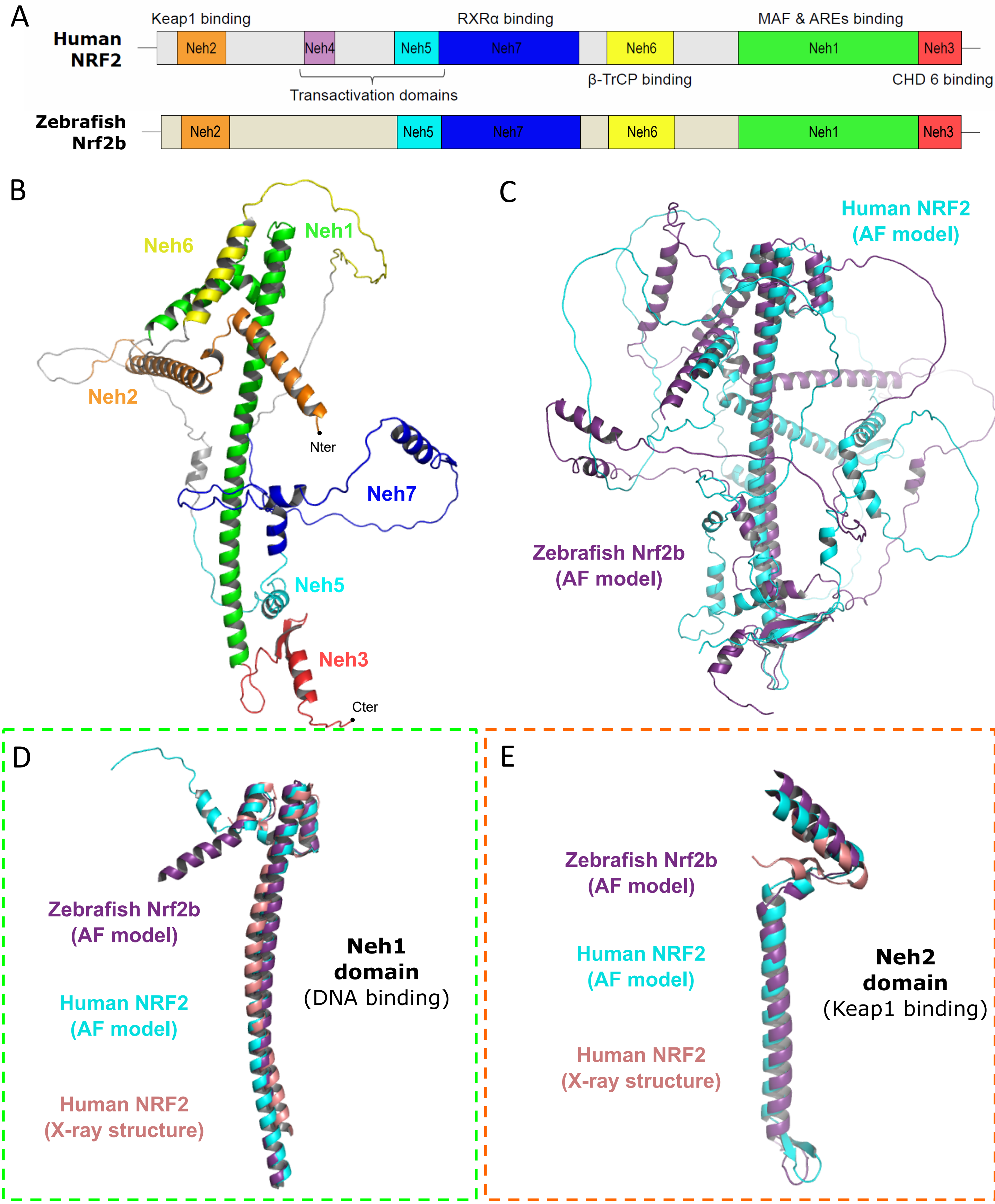
A
Human NRF2
Keap1 binding
RXRα binding
MAF & AREs binding
Neh2
Neh4
Neh5
Neh7
Neh6
Neh1
Neh3
Transactivation domains
β-TrCP binding
CHD 6 binding
Zebrafish Nrf2b
Neh2
Neh5
Neh7
Neh6
Neh1
Neh3
B
Neh6
Neh1
Neh2
Nter
Neh7
Neh5
Neh3
Cter
C
Human NRF2 (AF model)
Zebrafish Nrf2b (AF model)
D
Zebrafish Nrf2b (AF model)
Human NRF2 (AF model)
Human NRF2 (X-ray structure)
Neh1 domain (DNA binding)
E
Zebrafish Nrf2b (AF model)
Human NRF2 (AF model)
Human NRF2 (X-ray structure)
Neh2 domain (Keap1 binding)

S2

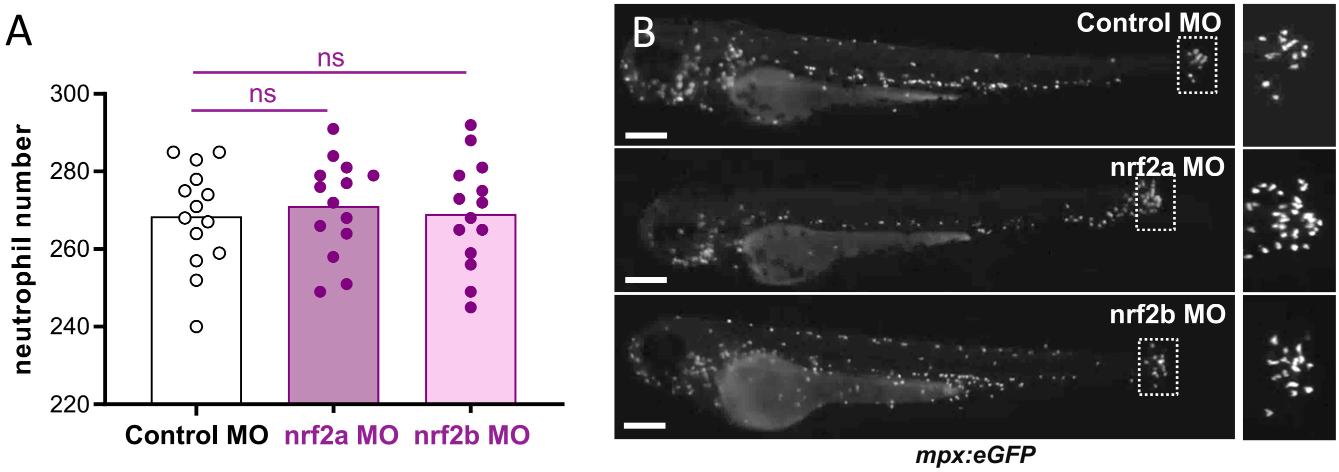
A
ns
ns
300
280
260
240
220
neutrophil number
Control MO
nrf2a MO
nrf2b MO
B
Control MO
nrf2a MO
nrf2b MO
mpx:eGFP

S3

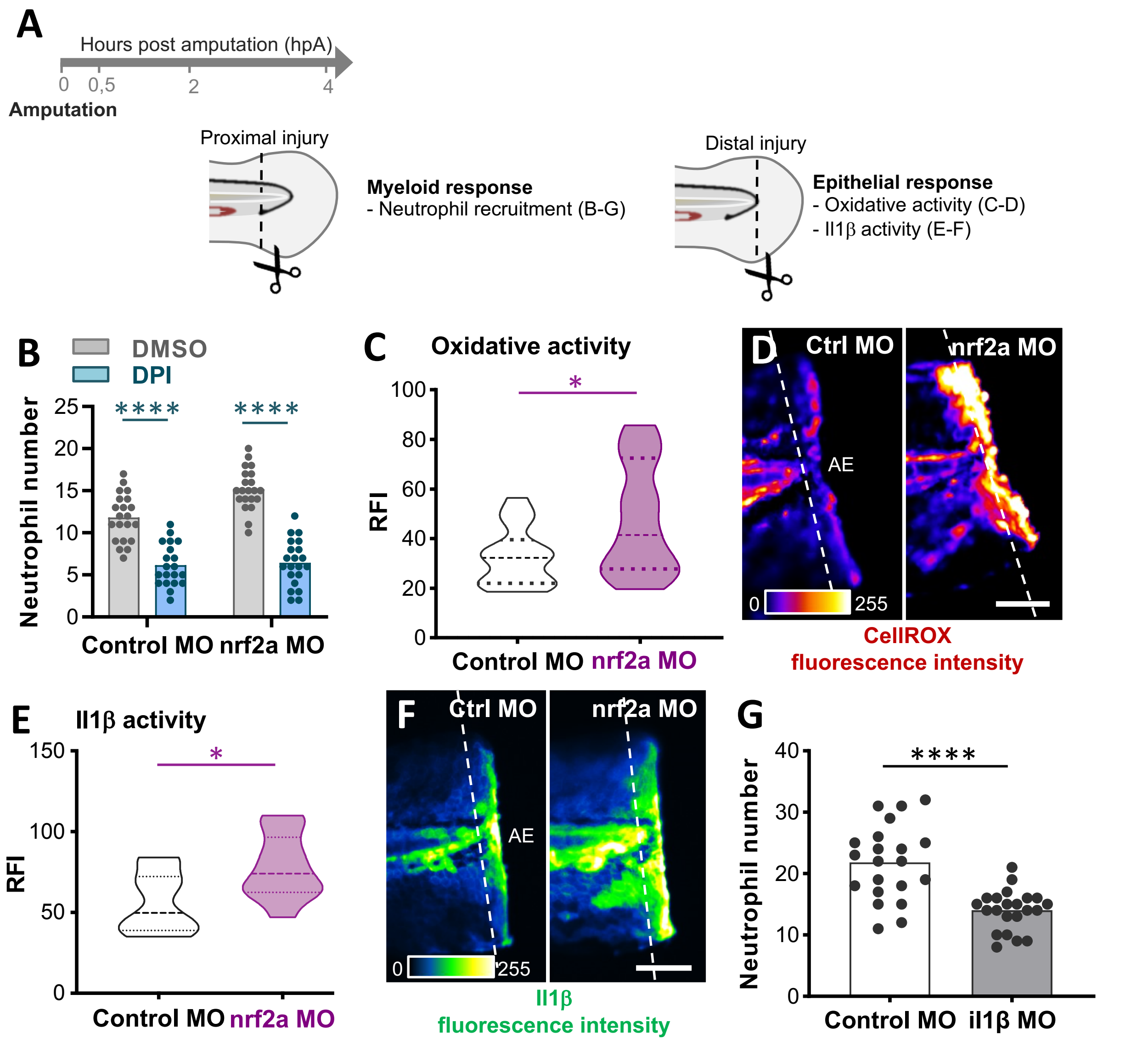
A
Hours post amputation (hpA)
0 0,5 2 4
Amputation
Proximal injury
Myeloid response
- Neutrophil recruitment (B-G)
Distal injury
Epithelial response
- Oxidative activity (C-D)
- Il1β activity (E-F)
B
DMSO
DPI
Neutrophil number
****
****
Control MO
nrf2a MO
C
Oxidative activity
RFI
*
Control MO
nrf2a MO
D
Ctrl MO
nrf2a MO
AE
0
255
CellROX fluorescence intensity
E
Il1β activity
RFI
*
Control MO
nrf2a MO
F
Ctrl MO
nrf2a MO
AE
0
255
Il1β fluorescence intensity
G
Neutrophil number
****
Control MO
il1β MO

S4

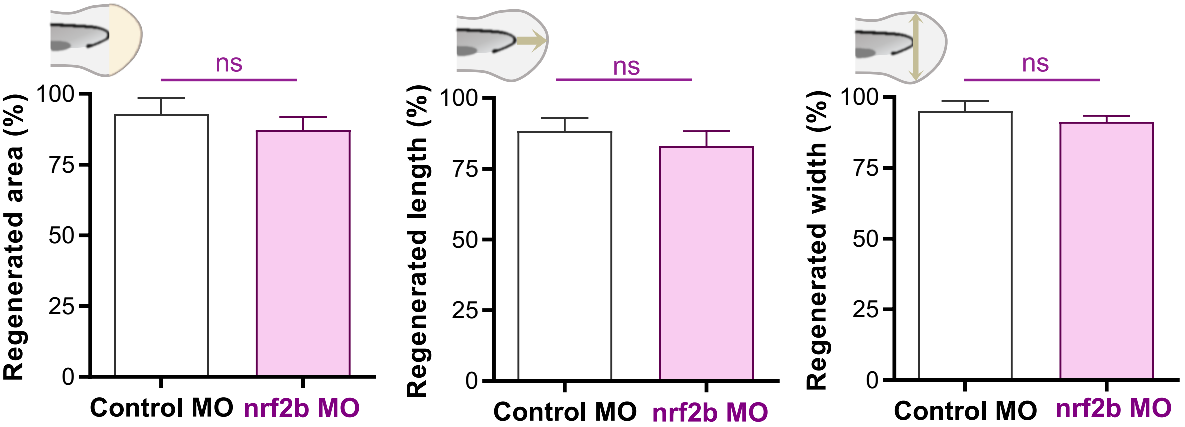
ns
ns
ns
Regenerated area (%)
Regenerated length (%)
Regenerated width (%)
100
75
50
25
0
Control MO
nrf2b MO

S5

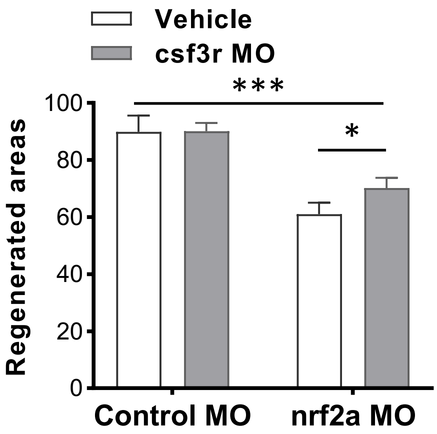
Vehicle
csf3r MO
***
*
Regenerated areas
100
80
60
40
20
0
Control MO
nrf2a MO

S6

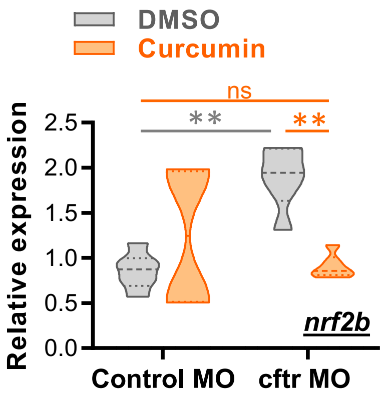

DMSO
Curcumin
ns
**
**
Relative expression
2.5
2.0
1.5
1.0
0.5
0.0
nrf2b
Control MO
cftr MO


S7

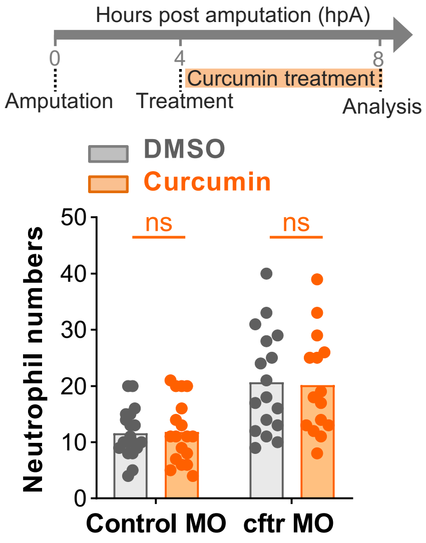

Hours post amputation (hpA)
0
4
8
Curcumin treatment
Amputation
Treatment
Analysis
DMSO
Curcumin
ns
ns
Neutrophil numbers
0
10
20
30
40
50
Control MO
cftr MO


S8

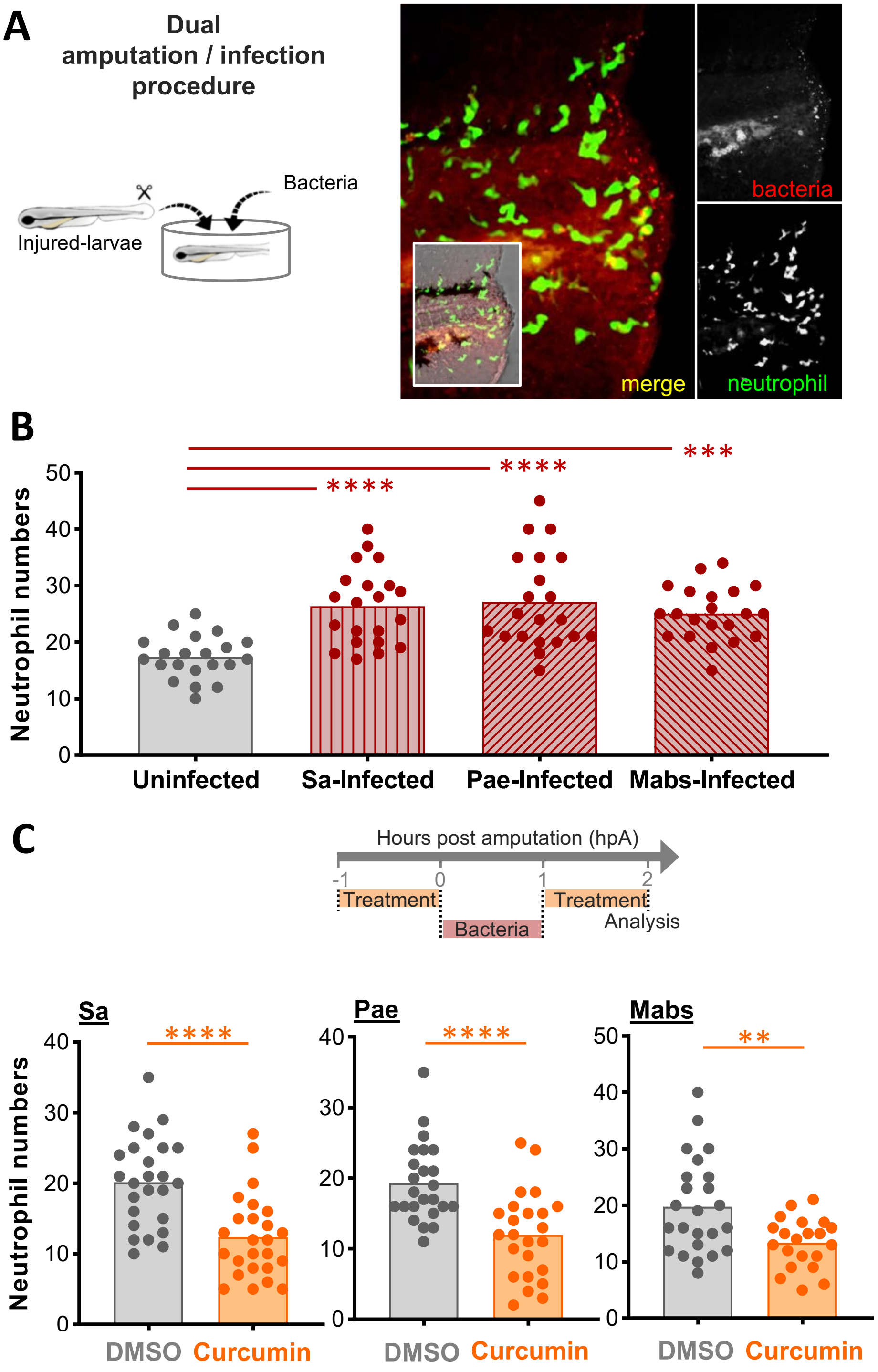
A
Dual amputation / infection procedure
Bacteria
Injured-larvae
bacteria
merge
neutrophil
B
Neutrophil numbers
****
****
***
Uninfected
Sa-Infected
Pae-Infected
Mabs-Infected
C
Hours post amputation (hpA)
-1
0
1
2
Treatment
Treatment
Bacteria
Analysis
Sa
Pae
Mabs
Neutrophil numbers
****
****
**
DMSO
Curcumin

S9

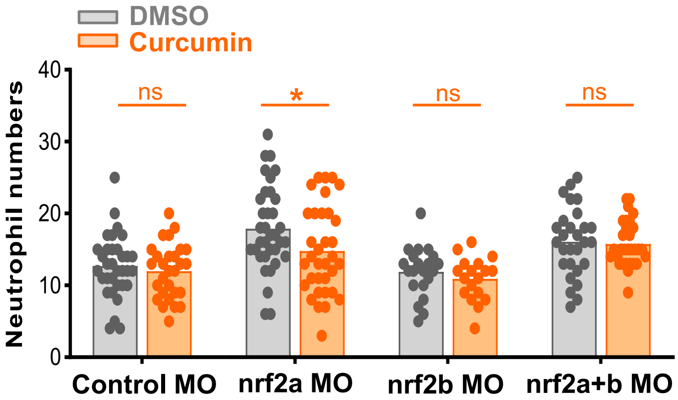
DMSO
Curcumin
Neutrophil numbers
40
30
20
10
0
ns
*
ns
ns
Control MO
nrf2a MO
nrf2b MO
nrf2a+b MO

S10

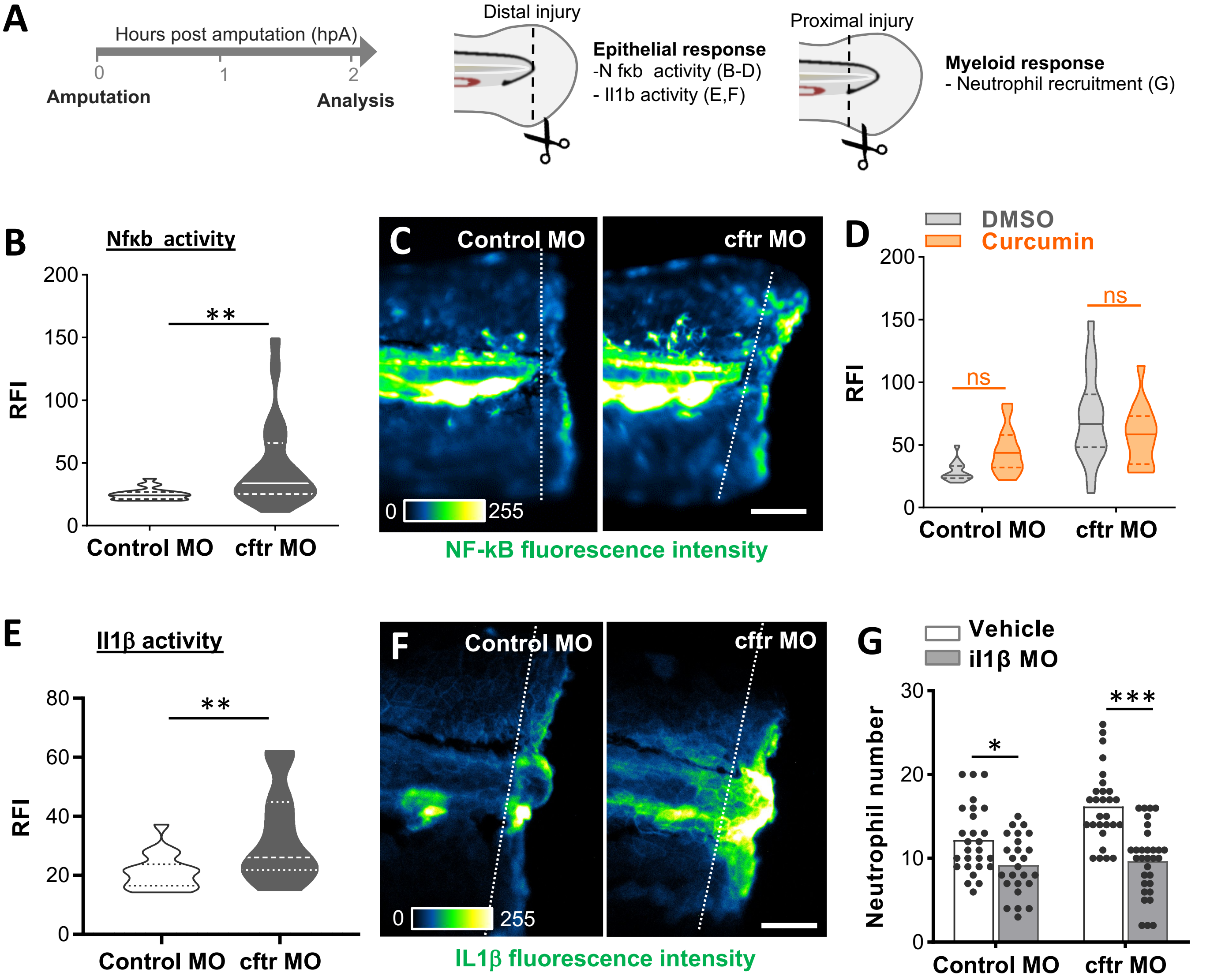
A
Hours post amputation (hpA)
0
1
2
Amputation
Analysis
Distal injury
Epithelial response
-N fκb activity (B-D)
- Il1b activity (E,F)
Proximal injury
Myeloid response
- Neutrophil recruitment (G)
B
Nfκb activity
RFI
**
Control MO
cftr MO
C
Control MO
cftr MO
0
255
NF-kB fluorescence intensity
D
DMSO
Curcumin
RFI
ns
ns
Control MO
cftr MO
E
Il1β activity
RFI
**
Control MO
cftr MO
F
Control MO
cftr MO
0
255
IL1β fluorescence intensity
G
Vehicle
il1β MO
Neutrophil number
*
***
Control MO
cftr MO

S11

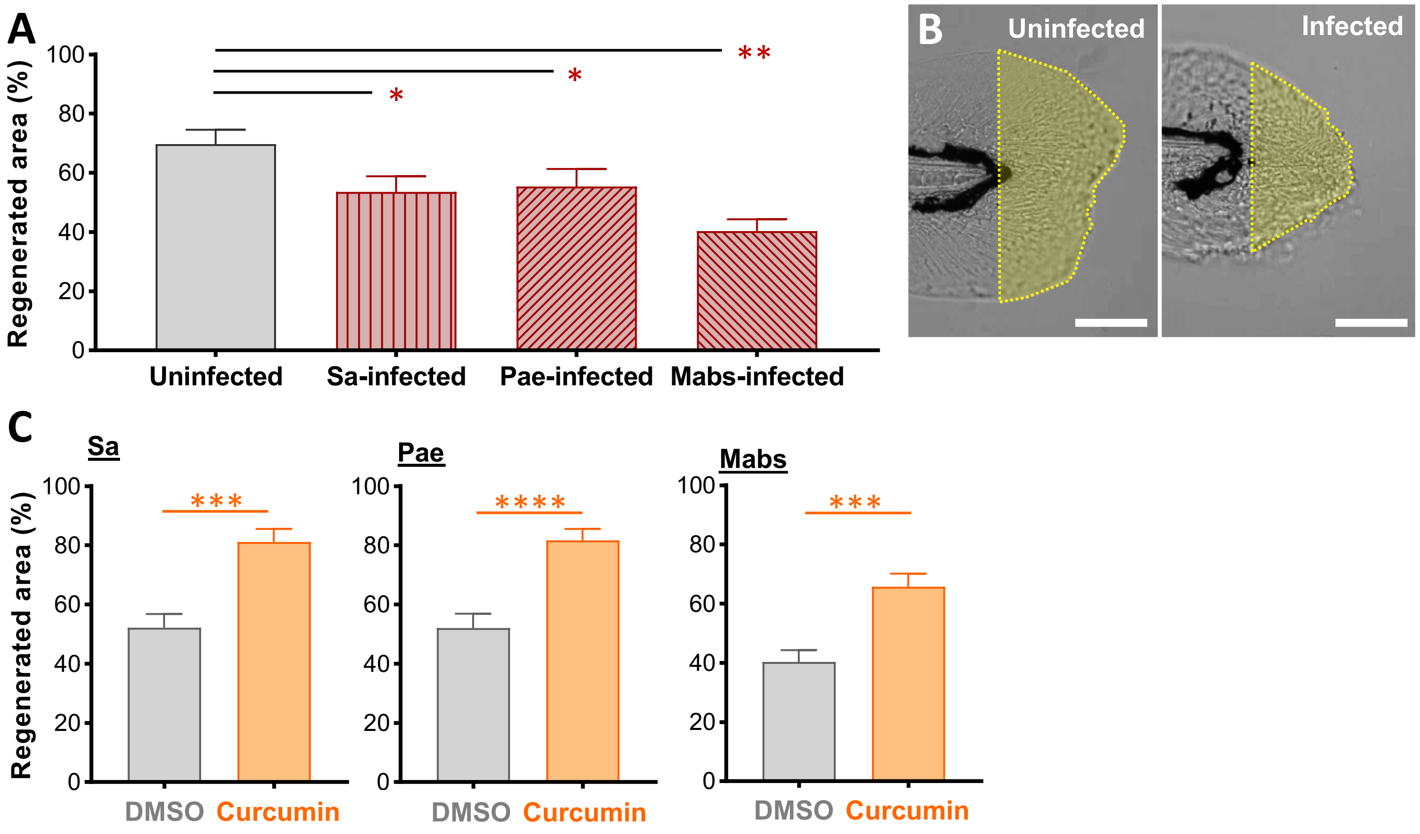
A
Regenerated area (%)
100
80
60
40
20
0
Uninfected
Sa-infected
Pae-infected
Mabs-infected
**
*
*
B
Uninfected
Infected
C
Sa
Pae
Mabs
***
****
***
DMSO
Curcumin

S12

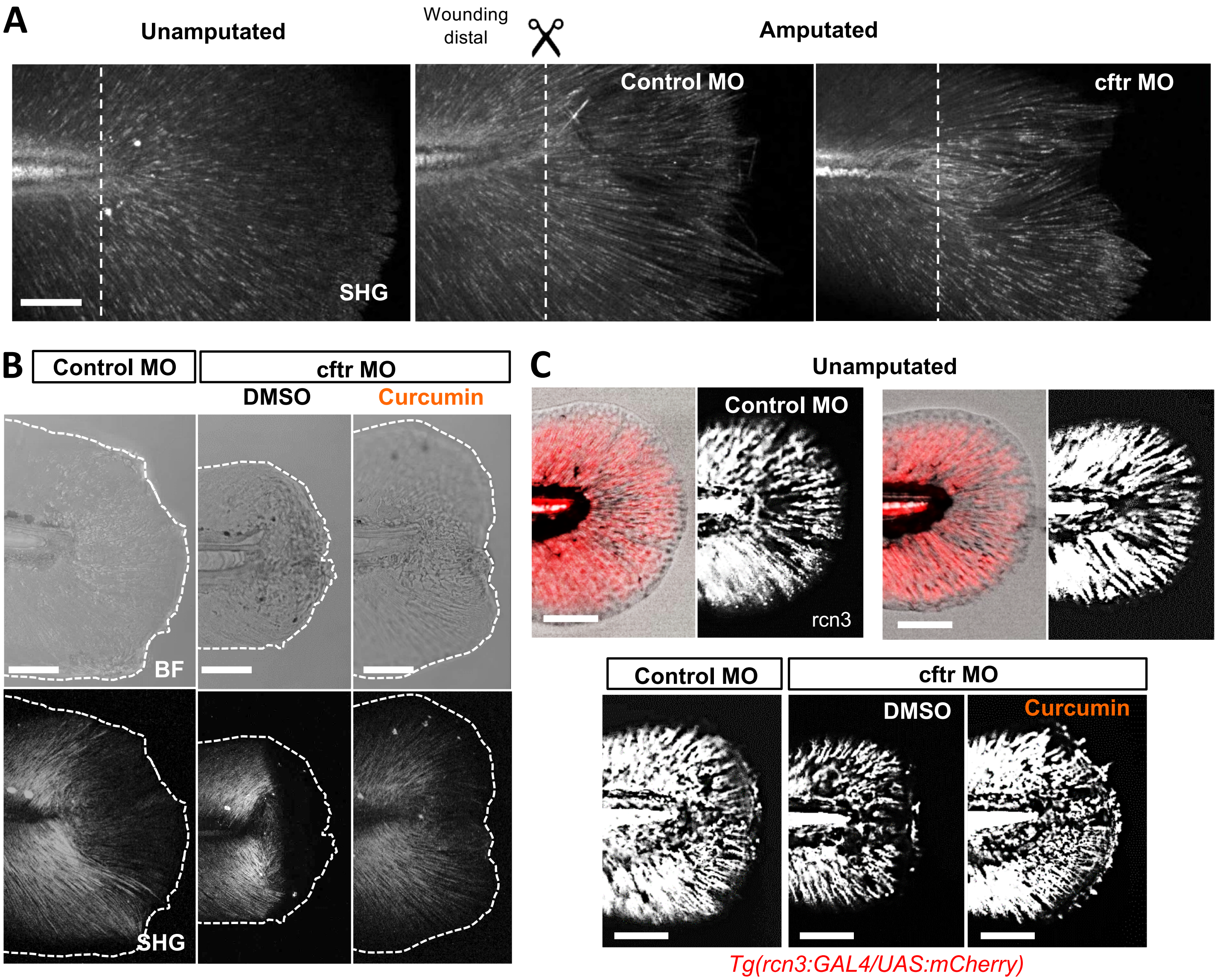

A
Unamputated
Wounding distal
Amputated
Control MO
cftr MO
SHG
B
Control MO
cftr MO
DMSO
Curcumin
BF
SHG
C
Unamputated
Control MO
rcn3
Control MO
cftr MO
DMSO
Curcumin
Tg(rcn3:GAL4/UAS:mCherry)


**S13**
